# Global increase in circRNA levels in myotonic dystrophy

**DOI:** 10.1101/489070

**Authors:** Karol Czubak, Katarzyna Taylor, Agnieszka Piasecka, Krzysztof Sobczak, Katarzyna Kozlowska, Anna Philips, Saam Sedehizadeh, J. David Brook, Marzena Wojciechowska, Piotr Kozlowski

## Abstract

Splicing aberrations induced as a consequence of the sequestration of MBNL splicing factors on the DMPK transcript, which contains expanded CUG repeats, present a major pathomechanism of myotonic dystrophy type 1 (DM1). As MBNLs may also be important factors involved in the biogenesis of circular RNAs (circRNAs), we hypothesized that the level of circRNAs would be decreased in DM1. To test this hypothesis, we selected twenty well-validated circRNAs and analyzed their levels in several experimental systems (e.g., cell lines, DM muscle tissues, and a mouse model of DM1) using droplet digital PCR assays. We also explored the global level of circRNAs using two RNA-Seq datasets of DM1 muscle samples. Contrary to our original hypothesis, our results consistently showed a global increase in circRNA levels in DM1 and we identified numerous circRNAs that were increased in DM1. We also identified many genes (including muscle-specific genes) giving rise to numerous (>10) circRNAs. Thus, this study is the first to show an increase in global circRNA levels in DM1. We also provided preliminary results showing the association of circRNA level with muscle weakness and alternative splicing changes that are biomarkers of DM1 severity.

**Author Summary:** Recently, a great deal of interest has been focused on a new class of RNA molecules called circular RNAs (circRNAs). To date, thousands of circRNAs have been found in different human cells/tissues. Although the function of circRNAs remains mostly unknown, circRNAs have emerged as an important component of the RNA-RNA and RNA-protein interactome. Thus, intensive efforts are being made to fully understand the biology and function of circRNAs, especially their role in human diseases. As an important role in the biogenesis of circRNA may be played by MBNL splicing factors, in this study we used DM1 (to a lesser extent, DM2) as a natural model in which the level of MBNLs is decreased. In contrast to the expected effect, our results consistently showed a global increase in circRNA levels in DM1. As a consequence, whole genome transcriptome analysis revealed dozens of circRNAs with significantly altered (mostly increased) levels in DM1. Furthermore, we observed that the circRNA levels were in many cases strongly associated with DM1 severity.

## INTRODUCTION

Myotonic dystrophy type 1 (*dystrophia myotonica 1*, DM1, OMIM: 160900) is the most common form of adult-onset muscular dystrophy, affecting approximately 1 in 8,000 people worldwide. DM1 is an autosomal dominant disorder caused by an expansion of CTG repeats in the 3′ untranslated region (3’UTR) of the *dystrophia myotonica protein kinase* (*DMPK*) gene (1-3). Unaffected individuals have between 5 and ~34 repeats, whereas in DM1 patients, the triplet repeat is expanded, often to hundreds or even thousands of copies (1). The pathogenesis of DM1 is strongly linked to the expression of mutation-containing transcripts and is manifested through the nuclear accumulation of mutant transcripts in characteristic foci (4). The presence of these mutant transcripts causes the sequestration of muscleblind-like (MBNL) proteins [including MBNL1, the main MBNL family protein in muscles (5, 6), MBNL2, and MBNL3], which normally regulate alternative splicing of pre-mRNAs encoding proteins critical for skeletal, cardiac, and nervous system function (7, 8). Thus, their sequestration and functional insufficiency result in aberrant alternative splicing of many target genes. For example, mis-splicing of the *CLCN1* exon 7, the *INSR* exon 11, and the *BIN1* exon 11 were shown to be associated with reduced chloride conductance, lower insulin responsiveness, and muscle weakness, respectively (9-13). A pathomechanism similar to that observed in DM1 was also proposed for myotonic dystrophy type 2 (*dystrophia myotonica 2*, DM2, OMIM: 602668), a disease caused by an expansion of CCTG repeats in the first intron of the *CCHC-type zinc finger nucleic acid binding protein* (*CNBP*) gene (14). However, in this study, we mainly focused on DM1.

The results of a recent study suggest that in addition to a function in alternative splicing, MBNLs may play an important role in the biogenesis of a recently recognized class of RNA molecules called circular RNAs (circRNAs) (15). Unlike other types of RNA, circRNAs are very stable molecules. Due to the low expression level of the initially identified circRNAs, they were considered byproducts of aberrant RNA splicing. However, with the dissemination of RNA-Seq technology, research has revealed that circRNAs are abundant among a variety of transcriptomes (16, 17). Although the levels of most circRNAs are low, there are examples of circRNAs with levels comparable to or higher than those of their linear counterparts (18). Most circRNAs are encoded by protein-coding genes and derived from their exons, which may indicate that transcription of circRNAs is directed by RNA polymerase II and that their biogenesis is mediated by the spliceosome. In the majority of cases, head-to-tail junctions of circular transcripts are flanked by canonical splice sites (15, 19). Reportedly, the formation of circRNAs may occur both posttranscriptionally and cotranscriptionally (15, 20, 21), and their biogenesis competes with the formation of linear transcripts (mRNA). The mechanisms of this competition are tissue-specific and conserved from flies to humans (15, 22). To date, no function has been assigned for the vast majority of circRNAs, with exceptions such as circCDR1as, *Sry* circRNA or circHIPK3 (hsa_circ_0000284), which can act as microRNA sponges (17, 23, 24). Other functions, such as involvement in protein and/or RNA transport (17), regulating synaptic functions in neural tissue (25) or acting as templates for translation of functional peptides [e.g., (26)], have also been proposed for circRNAs.

The precise mechanism of circRNA generation remains unknown. However, several mechanisms of circRNA biogenesis have been proposed (16, 18, 27). All of these proposed mechanisms assume the generation of circRNAs by head-to-tail splicing (back-splicing). One of the proposed mechanisms suggests that RNA-binding proteins (RBPs), which bind to specific motifs in introns flanking circRNA-coding exons, play an important role in circRNA biogenesis (15, 28). Back-splicing is facilitated by the interaction between RBPs, which bring the introns closer together. The *Drosophila* Mbl protein (orthologue of human MBNLs) may be a circRNA-biogenesis RBP (15). Interestingly, one of Mbl-regulated circRNAs is circMBNL1/circMbl, a circRNA generated from the second exon of the *MBNL1/Mbl* gene. The introns flanking this circRNA contain highly conserved MBNL/Mbl-binding motifs. Furthermore, the exogenous expression of Mbl stimulates circRNA production from endogenous MBNL1/Mbl transcripts in both humans and flies. Mbl-binding sequences in both introns are necessary, suggesting that Mbl induces circularization by bridging the two flanking introns. Importantly, downregulation of Mbl in both fly cell culture and fly neural tissue leads to a significant decrease in circMbl level whereas the elevated level of Mbl increases the level of circMbl as well as other circRNAs, suggesting a general role for MBNLs/Mbl in circRNA biogenesis (15).

In this work, we aimed to understand the link between nuclear sequestration of MBNL proteins and the expression levels of circRNAs. Since MBNL proteins may be involved in circRNA biogenesis (15), we hypothesized that the generation of circRNAs would be downregulated by the diminished functional levels of MBNLs, which are sequestered in mutant RNA foci (7). To test this hypothesis, we selected twenty well-validated circRNAs and analyzed their expression levels in several experimental systems, including cultured human myoblasts and skeletal muscle biopsy samples from patients and healthy individuals. In addition, we used muscles from the *HSA*^LR^ transgenic mouse model of DM1 (29, 30) and human muscle cell lines with depletion of MBNL1 or MBNL3 (31). The analysis of circRNA expression levels was performed with in-house-designed droplet-digital PCR (ddPCR) (32, 33) assays. We also expanded this analysis and explored global levels of circRNAs using RNA-Seq data from an “exploratory cohort” of DM1 muscle samples of quadriceps femoris (QF) and tibialis anterior (TA) (http://www.dmseq.org/).

In summary, we found no downregulation of the analyzed circRNAs in DM (both DM1 and DM2) samples compared to those in non-DM samples. Therefore, these results question the role of MBNL proteins in circRNA biogenesis in muscles. Interestingly, in our experimental systems which are characterized by a lower level of functional MBNLs, we discovered a consistent increase in circRNA levels. As a result, we identified a subset of circRNAs that were upregulated in DM1 samples and could be used as novel biomarkers. Although the obtained data do not confirm our hypothesis regarding the link between MBNL sequestration and disrupted circRNA biogenesis in DM1 (and DM2), we do not exclude the possibility of the existence of individual circRNAs that are regulated by MBNLs. Additionally, we demonstrated that elevated circRNA levels associate with molecular (alternative splicing) and clinical (muscle weakness) symptoms of DM severity. However, the role of individual circRNAs altered in DM1 and their global function in DM1 pathogenesis remain to be determined.

## RESULTS

### Selection of circRNA species for expression analysis in DM1

To check whether the level of individual circRNAs is affected in DM1, we selected twenty circRNAs reported in previous studies (16-18, 22, 34) and deposited in circBase [(35); http://www.circbase.org/]. To avoid falsely identified circRNAs, we considered only circRNAs validated by at least 20 NGS reads in at least 2 previous studies. Fourteen circRNAs (Table 1) were selected based on their relatively high levels (compared to other circRNAs) in different types of cells/tissues and relatively high [≥10% in (18)] expression levels compared to that of their linear counterparts (mRNAs). Four circRNAs (Table 1) were selected based on a high number (≥10) of potential MBNL-binding sites [YGCY motifs; (36)] in adjacent (300 nt upstream and 300 nt downstream) sequences of their flanking introns. Additionally, we selected circCDR1as (hsa_circ_0001946) (17, 23), the well-studied circRNA generated from the antisense transcript of the *CDR1* gene (CDR1as), and circMBNL1 (hsa_circ_0001348) (Table 1), which derives from the second exon of *MBNL1*, which is reportedly involved in the self-regulation of *MBNL1* expression (37) and linked to circRNA biogenesis (15).

**Table 1.** CircRNAs selected for analysis.

| circRNA | circBase ID | genome localization<br>(hg 19) | homing<br>gene | circRNA:mRNA<br>ratio (18) | number of<br>potential<br>MBNL-binding<br>motifs |
| --- | --- | --- | --- | --- | --- |
| **circRNAs selected based on high level in different cells/tissues** |  |  |  |  |  |
| circASXL1 | hsa_circ_0001136 | 20:30954186\|30956926 | ASXL1 | 286% | 6 |
| circCASMAP1 | hsa_circ_0001900 | 9:138773478\|138774924 | CASMAP1 | 253% | 10 |
| circFAM13B | hsa_circ_0001535 | 5:137320945\|137324004 | FAM13B | 34% | 5 |
| circHIPK3 | hsa_circ_0000284 | 11:33307958\|33309057 | HIPK3 | 721% | 2 |
| circMBOAT2 | hsa_circ_0000972 | 2:9048750\|9098771 | MBOAT2 | 19% | 5 |
| circMIB1 | hsa_circ_0000835 | 18:19345732\|19359646 | MIB1 | 26% | 1 |
| circNFATC3 | hsa_circ_0000711 | 16:68155889\|68160513 | NFATC3 | 52% | 3 |
| circPHC3 | hsa_circ_0001359 | 3:169854206\|169867032 | PHC3 | 22% | 5 |
| circPIP5K1C | hsa_circ_0000871 | 19:3660963\|3661999 | PIP5K1C | 11% | 2 |
| circSCMH1 | hsa_circ_0000061 | 1:41536266\|41541123 | SCMH1 | 23% | 3 |
| circSHKBP1 | hsa_circ_0000936 | 19:41089303\|41089623 | SHKBP1 | 14% | 9 |
| circUBAP2_e7-8 | hsa_circ_0001851 | 9:33971648\|33973235 | UBAP2 | 82% | 4 |
| circUBAP2_e9-12 | hsa_circ_0001847 | 9:33953282\|33963789 | UBAP2 | 63% | 3 |
| circZKSCAN1 | hsa_circ_0001727 | 7:99621041\|99621930 | ZKSCAN1 | 99% | 14 |
| **circRNAs selected based on high number of potential MBNL-binding motifs** |  |  |  |  |  |
| circCCDC134 | hsa_circ_0001238 | 22:42204878\|42206295 | CCDC134 | 98% | 10 |
| circFOXK2 | hsa_circ_0000816 | 17:80521229\|80526077 | FOXK2 | 26% | 12 |
| circPDCD11 | hsa_circ_0000258 | 10:105197771\|105198565 | PDCD11 | 129% | 11 |
| circPROSC | hsa_circ_0001788 | 8:37623043\|37623873 | PROSC | 15% | 11 |
| **circRNAs additionally included in the analysis** |  |  |  |  |  |
| circCDR1as | hsa_circ_0001946 | X:139865339\|139866824 | CDR1as | - | 5 |
| circMBNL1 | hsa_circ_0001348 | 3:152017193\|152018156 | MBNL1 | - | 7 |

### Design of assays to analyze circRNA expression

For each selected circRNA, we designed PCR assays allowing amplification and parallel analysis of a given circRNA and its linear mRNA counterpart. Each assay consisted of three primers as follows: one primer common to both the circular and linear transcript and two primers specific for either the circular or linear transcript (Figure 1A, Table S1). The size of circRNA-specific amplicons was analyzed by agarose gel electrophoresis (Figure 1B), and the predicted back-splice sites were subsequently confirmed by Sanger sequencing (Figure 1C and Figure S2). The specific assays were employed for quantification of cDNA copies corresponding to circRNA and linear mRNA transcripts using ddPCR that enables absolute quantification of nucleic acid templates (32, 33). ddPCR involves partitioning the analyzed sample into many low-volume droplet reactions, and only a fraction of these reactions contains one (in most cases) or more template molecules (positive droplets). The final concentration of the analyzed templates was determined by Poisson statistical analysis of the number of positive and negative droplets. The level of a given circRNA in each analyzed sample was calculated as fraction of circular particles (FCP; see Materials and methods). Next, the FCPs calculated for DM1 and control samples were compared (Figure 1D, E).

**Figure 1.**
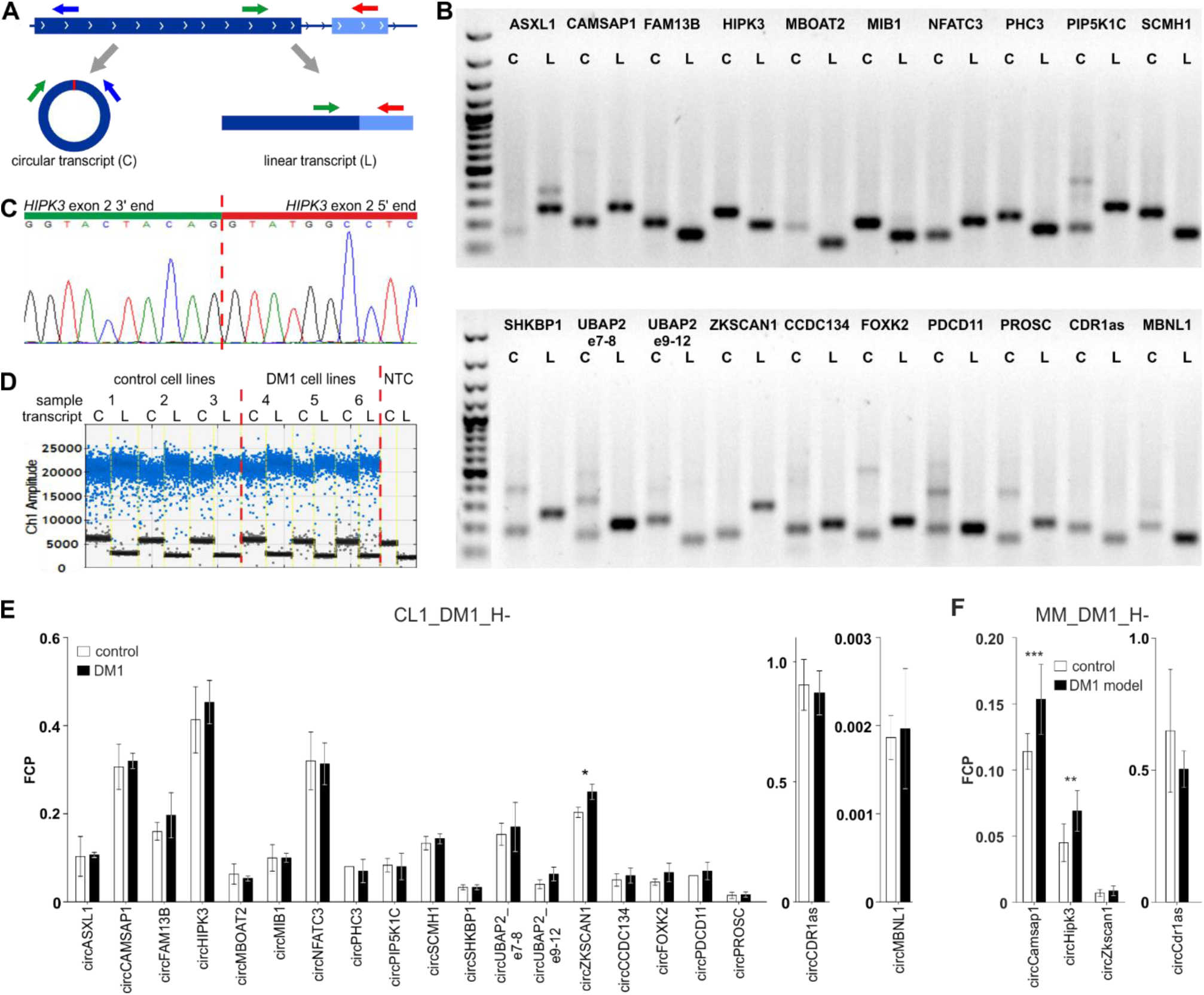
Design and validation of the assays used for analysis of circRNA levels. **A)** Each assay consisted of three primers as follows: one primer (green arrow) common to the circular (C) and linear (L) transcripts (circRNA and mRNA, respectively) and two primers specific for either circular (blue arrow) or linear (red arrow) transcripts. The primers specific to linear transcripts were located in either the downstream or upstream exon, outside of circRNA-coding exons. **B)** Gel electrophoresis confirming the size of circRNA-specific (C) and linear, mRNA-specific (L) amplicons. Additional bands in some tracks corresponding to circRNAs indicate the occurrence of circRNA-related concatemers [see (25)]. An additional band in *ASXL1* linear transcript track corresponds to the alternative transcript containing alternatively included (97-nt long) exon 5. The first track is the GeneRuler 1 kb DNA Ladder (Thermo Fisher Scientific, Waltham, MA, USA) **C)** Exemplary result of Sanger sequencing of the predicted back-splice site of circHIPK3. Results of sequencing of back-splice sites of other circRNAs are shown in Figure S2. **D)** Exemplary result of the ddPCR analysis of circHIPK3 in the myoblast cell line (CL1_DM1_H-) sample set. Sample number and type are indicated above the graph. NTC–no template control. Ch1 Amplitude–relative fluorescence signal in channel 1. Each blue dot represents one copy of either circular or linear transcript (positive droplets), while the black dots represent negative (empty) droplets. For each sample, the number of positive and negative droplets was used to calculate the concentration of the analyzed transcript. Next, for each cell line, the level of circular transcripts was calculated as FCP. **E)** Bar graph showing the levels of circRNAs (expressed as FCPs) in control (white bars) and DM1 (black bars) myoblast cell line samples (CL1_DM1_H-). The values indicated by white or black bars are the averaged FCPs calculated for either three control or three DM1 cell lines (shown in D). The whiskers represent standard deviation (SD) values. The differences in circRNA levels were compared with t-tests, and p-values <0.05 are indicated by asterisks. **F)** Bar graph showing the level of circRNAs analyzed in the *HSA*^LR^ transgenic mouse model of DM1 and in control background (*FVB*) mice (MM_DM1_H-sample set). The scheme of bar graph is as in Figure 1E. Asterisks indicate the following significance level: ** – p<0,01; *** – p<0,001.

Additionally, gel electrophoresis of the PCR product specific for circMBNL1 revealed an additional longer band. Analysis of this additional band led to the identification and characterization of a new circRNA (circMBNL1’) consisting of the second exon of *MBNL1* and a 93-nt fragment of the large (~114 kb long) downstream intron 2 (Figure S3). The analysis of the surrounding sequence with the GENESCAN online tool (http://genes.mit.edu/GENSCAN.html) identified (with high confidence) the incorporated fragment of intron as an exon, with canonical 5’ and 3’ splice sites.

### Analysis of expression levels of the selected circRNAs in DM samples

Human myoblast cell lines (CL), as well as skeletal muscle biopsy (BP) tissues from DM1, DM2, and non-DM controls, were used to compare expression levels of circRNAs in DM and unaffected samples in 6 different sample sets (defined in Materials and methods). As shown in Figure 1E and Figure S4, none of the tested circRNAs were substantially decreased in either DM1 or DM2 samples (no matter of the used sample type or condition of reverse transcription). The marginally significant differences of the individual cicRNA levels are indicated by asterisks on the graphs. In contrast, a summary of the experiments performed with the use of either cell lines or patient-derived muscle samples showed that the selected circRNAs tend to be rather increased than decreased in DM samples (for summary, see Table 2 and Table S2). For example, in the muscle biopsy sample set (BP1_DM1_H-), 12 out of 15 tested circRNAs were increased in DM1 (Chi^2^, p-value=0.02). A similar effect was observed when the circRNA level was normalized against the levels of housekeeping genes (*GAPDH* and *ACTB;* data not shown).

**Table 2.**
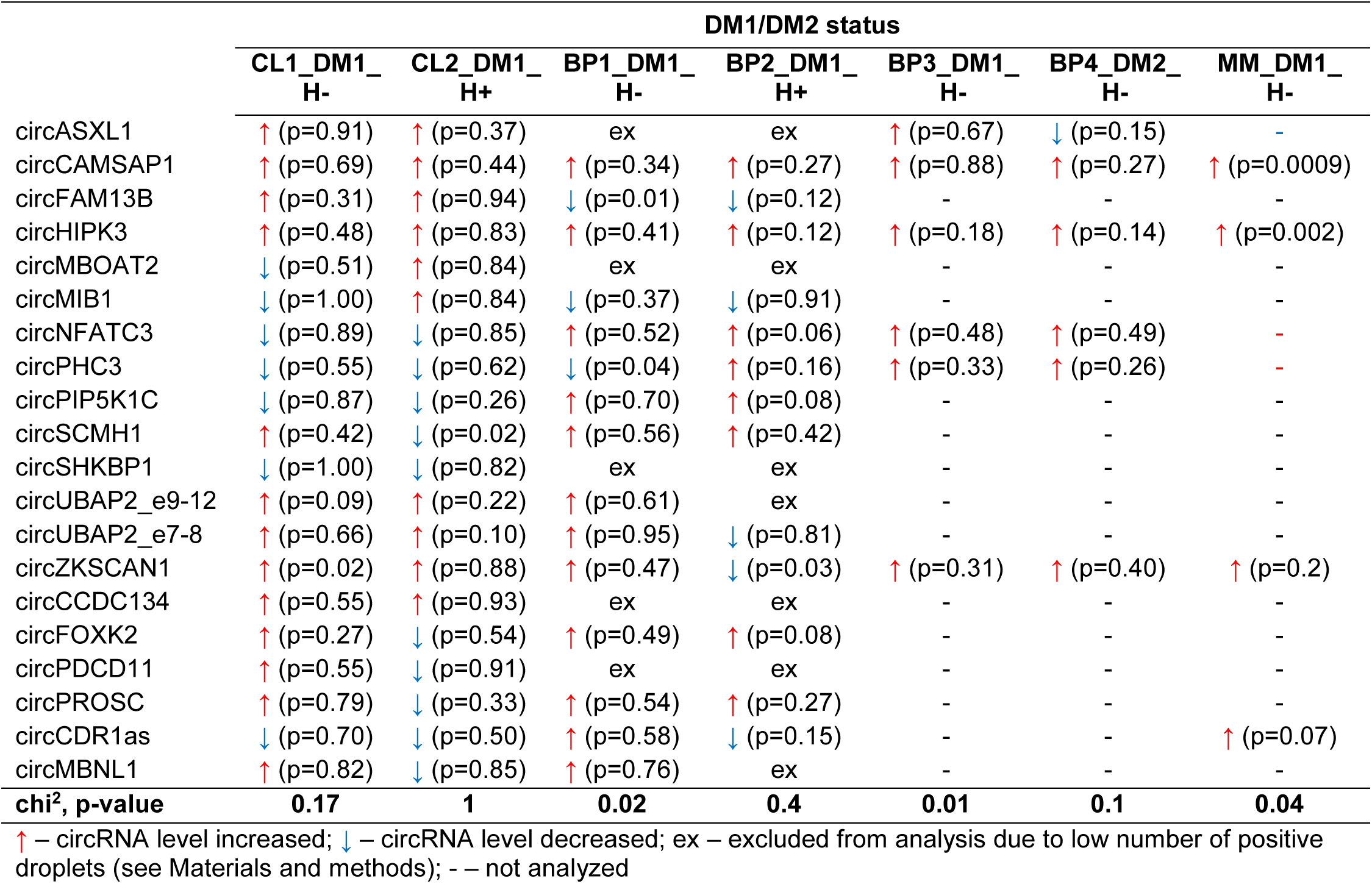
Results of experimental analyses of circRNA expression levels.

The disadvantage of analysis of human biopsy samples is that they may not always be of homogenous quality (e.g., different sample sources or divergent tissue and/or RNA treatment protocols may result in differences in RNA integrity). Moreover, the limited access to this type of samples and consequently small sample sets does not always allow the detection (with appropriate statistical support) of smaller changes in the levels of analyzed transcripts. Therefore, in the next step, we used cDNA samples from muscles of the commonly used and well-characterized mouse model of DM1 [*HSA*^LR^, (29)] and compared them with samples from control background (*FVB*) mice. For analysis, we selected four mouse circRNAs (circCamsap1, circHipk3, circZkscan1, and circCdr1as) that are orthologues of the human circRNAs analyzed in this study. As shown in Figure 1F, the levels of two circRNAs (i.e., circCamsap1 and circHipk3) were significantly increased in *HSA*^LR^ (t-test p=0.001, and p=0.002, respectively).

In conclusion, our experimental analyses do not support our original hypothesis, assuming a decrease in the circRNA level in DM. Additionally, some of the results suggest rather the opposite effect, i.e. a trend toward increase of circRNA level in DM (although in most cases only marginally significant).

### Analysis of circRNA levels in DM1 with RNA-Seq datasets

CircRNAs selected for the experiments described above may not be representative, and global circRNA level changes may be too small to be detected with a few circRNAs. Therefore, in the next step, to better evaluate the global circRNA level, we used the RNA-Seq data deposited in the DMseq database [(35); http://www.dmseq.org/]. For the analysis, we selected data sets of muscle samples most frequently represented in the database, QF muscle (11 control samples and 12 DM1 samples) and TA muscle (6 control samples and 21 DM1 samples). To avoid potential technical variations in analysis, we selected only samples with sequencing data generated with uniform procedures (for details, see Materials and methods). For the detection and quantification of circRNAs and their linear mRNA counterparts, we used one of the existing computational circRNA-detection tools, CIRI2 (38), which uses maximum likelihood estimation based on multiple seed matching. This tool enables the identification of back-spliced junction reads and the filtration of false positives derived from repetitive sequences and mapping errors.

In total, in QF samples, we detected 22,816 distinct circRNAs (‘all’; a substantial fraction were confirmed by just a few reads), 4,168 (18%) of which were classified as ‘validated’ (confirmed by at least five reads in at least two samples), and 152 (0.7%) were classified as ‘common’ (present in all or all but one sample of either control or DM1 samples). In the case of TA samples, the ‘all’ group contained 38,403 circRNAs, and the ‘validated’ and ‘common’ groups contained 7,537 (20% of ‘all’) and 403 (1% of ‘all’) circRNAs, respectively. As expected, the fraction of known [deposited in circBase and in (39)] circRNAs increased with the level of validation in both QF and TA (Table 3, Table S3, Table S4).

**Table 3.**
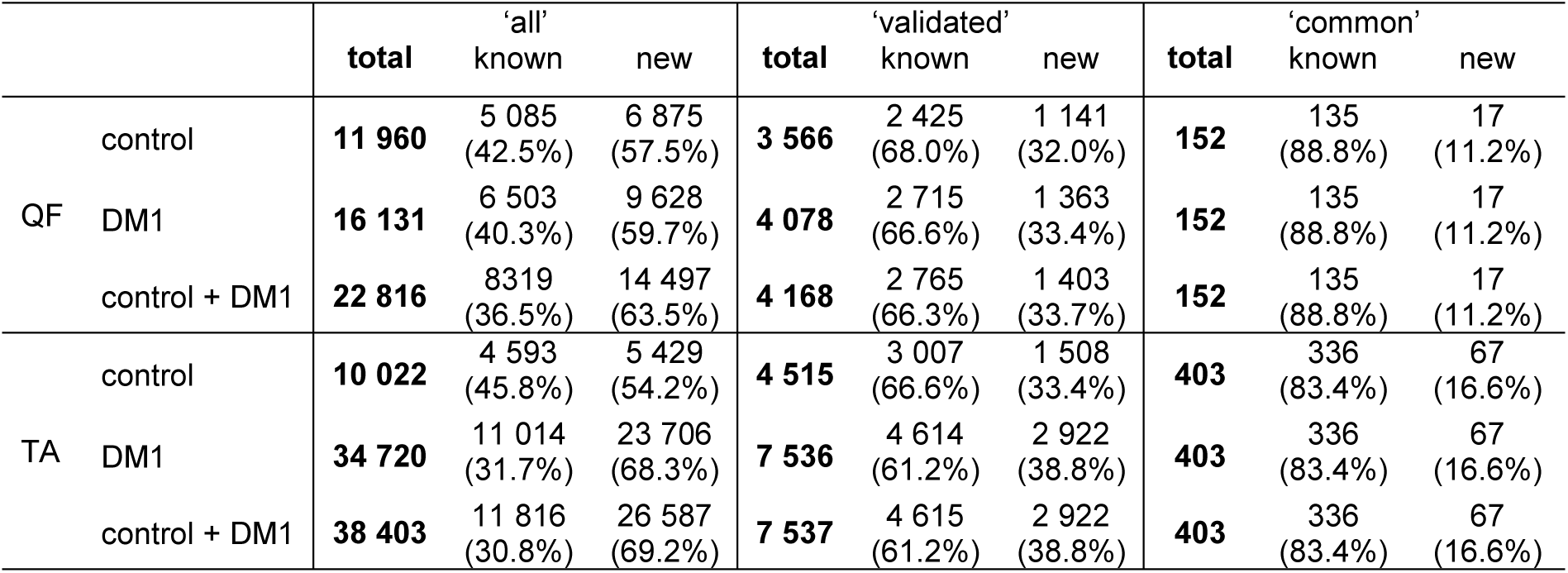
Number of circRNAs in QF and TA tissues in different validation groups.

|  |  | total | 'all'<br>known | new | total | 'validated'<br>known | new | total | 'common'<br>known | new |
| --- | --- | --- | --- | --- | --- | --- | --- | --- | --- | --- |
| QF | control | **11 960** | 5 085<br>(42.5%) | 6 875<br>(57.5%) | **3 566** | 2 425<br>(68.0%) | 1 141<br>(32.0%) | **152** | 135<br>(88.8%) | 17<br>(11.2%) |
|  | DM1 | **16 131** | 6 503<br>(40.3%) | 9 628<br>(59.7%) | **4 078** | 2 715<br>(66.6%) | 1 363<br>(33.4%) | **152** | 135<br>(88.8%) | 17<br>(11.2%) |
|  | control + DM1 | **22 816** | 8319<br>(36.5%) | 14 497<br>(63.5%) | **4 168** | 2 765<br>(66.3%) | 1 403<br>(33.7%) | **152** | 135<br>(88.8%) | 17<br>(11.2%) |
| TA | control | **10 022** | 4 593<br>(45.8%) | 5 429<br>(54.2%) | **4 515** | 3 007<br>(66.6%) | 1 508<br>(33.4%) | **403** | 336<br>(83.4%) | 67<br>(16.6%) |
|  | DM1 | **34 720** | 11 014<br>(31.7%) | 23 706<br>(68.3%) | **7 536** | 4 614<br>(61.2%) | 2 922<br>(38.8%) | **403** | 336<br>(83.4%) | 67<br>(16.6%) |
|  | control + DM1 | **38 403** | 11 816<br>(30.8%) | 26 587<br>(69.2%) | **7 537** | 4 615<br>(61.2%) | 2 922<br>(38.8%) | **403** | 336<br>(83.4%) | 67<br>(16.6%) |

To compare the global level of circRNA in control and DM1 samples, in each sample we summarized the number of reads [normalized as reads per million mappable reads (RPMs)] mapping to back-splice sequences (circRNA level) and mapping to the corresponding linear-splice sequences (linear mRNA level). As shown in Figure 2, the average global level of ‘all’ circRNAs was significantly increased in DM1 samples (p=0.002 in QF and p<0.0001 in TA). Importantly, no difference was detected compared with corresponding linear transcripts (p=0.5 and p=0.2 in QF and TA, respectively). The increased level of circRNA in DM1 samples was also visible for ‘validated’ and ‘common’ circRNAs (Figure S5). Similar results were obtained when the level of transcripts (number of reads) was normalized against the level of individual housekeeping genes, e.g., *ACTB* or *GAPDH* (data not shown).

**Figure 2.**
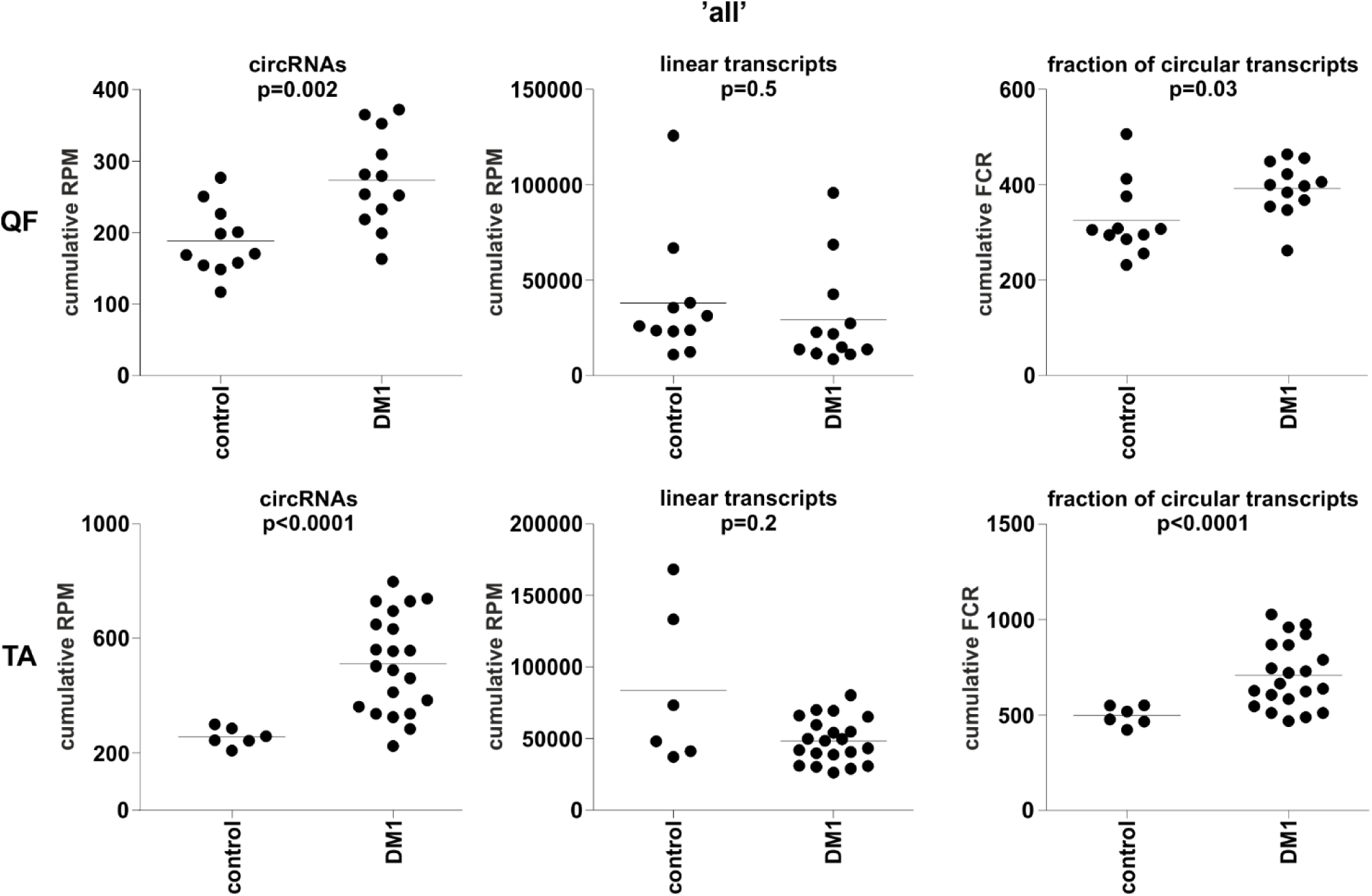
Comparison of cumulative levels of circRNAs and linear RNAs in control and DM1 muscle samples. Dot plots depicting the cumulative level of ‘all’ circRNAs and linear transcripts in QF (upper panel) and TA (lower panel). From the left (in each panel): cumulative RPM of circRNAs, cumulative RPM of linear transcripts, and cumulative FCR. The FDR-corrected p-value (t-test with correction for not-equal variance) of the differences between control and DM1 samples is shown above each dot plot.

The above changes in circRNA levels may be a reflection of an increase or decrease of expression from a particular gene or genome region. To control for this effect, we also normalized the levels of circRNAs against the levels of their linear counterparts, calculating the level of circRNAs as fraction of circRNA-specific reads in a total number of circRNA-specific and corresponding linear reads (FCR). Again, the cumulative value or averaged FCRs were higher in DM1 samples than in control samples (right graphs in Figure 2 and Figure S5). Additionally, in this analysis, circular transcripts of ‘common’ circRNAs accounted for ~5-10% of their linear counterparts.

### Differential expression of individual circRNAs

Although it was not the main purpose of the study, by using the generated data, we also analyzed the differential expression of individual circRNAs. This analysis was limited to only the sets of ‘common’ circRNAs (n=152 in QF and n=403 in TA) with expression levels detectable in the vast majority of analyzed samples. The difference in circRNA levels was calculated for the level of circRNAs normalized as RPMs and FCRs of individual circRNAs and expressed as log2 of fold change in DM1 samples vs. control samples. In both QF and TA, the changes in circRNA levels calculated with two normalization methods were highly correlated (Figure S6), indicating that circRNA changes do not depend on the expression of genes (level of their primary transcripts) from which they are generated. The results of the analyses are shown in Table S5 and Table S6 and graphically summarized in the form of volcano plots (Figure 3). The lists of the top ten most highly differentiated circRNAs (with respect to their RPM value) in QF and TA are shown in Table 4. As shown in Figure 3, log2 fold change values are substantially shifted toward positive values, indicating an excess of circRNAs with increased levels in DM1 samples. This effect is in line with the global increase in circRNA levels in DM1 (in both QF and TA) described above. For example, assuming that results fulfilling the following thresholds are significant (p-value<0.05 and log2 fold change ≤-1 or ≥1), we obtained 38 and 120 differentially expressed circRNAs in QF and TA, respectively. Among these circRNAs, circRNAs with increased expression levels in DM1 (Figure 3) were substantially overrepresented [i.e., 36 (95%) in QF (chi^2^, p<0.0001) and 104 (87%) in TA (chi^2^, p<0.0001)]. Similar bias toward circRNAs increased in DM1 may also be seen with other methods of normalization (e.g., such as FCR or normalization against the level of housekeeping genes; data not shown) as well as with other cutoff thresholds. Among circRNAs for which both RPM and FCR values were decreased in DM1 we studied whether MBNL may contribute to their biogenesis. We conducted the analysis of introns (300 nt upstream and 300 nt downstream from circRNA-generating exons) flanking these circRNAs. However, we did not show enrichment of potential MBNL-binding motifs (n ranging from 1 to 9, in most cases n≤5) that would justify the role of MBNLs in their biogenesis. The only interesting exception was circGSE1 (having as many as 29 potential MBNL-binding sites), with decreased RPM and RCF values in DM1 in TA (log2 fold change=-2.1; false discovery ratio (FDR)-corrected p-value=0.0001 and log2 fold change=-0.9; FDR-corrected p-value=0.1, respectively).

**Figure 3.**
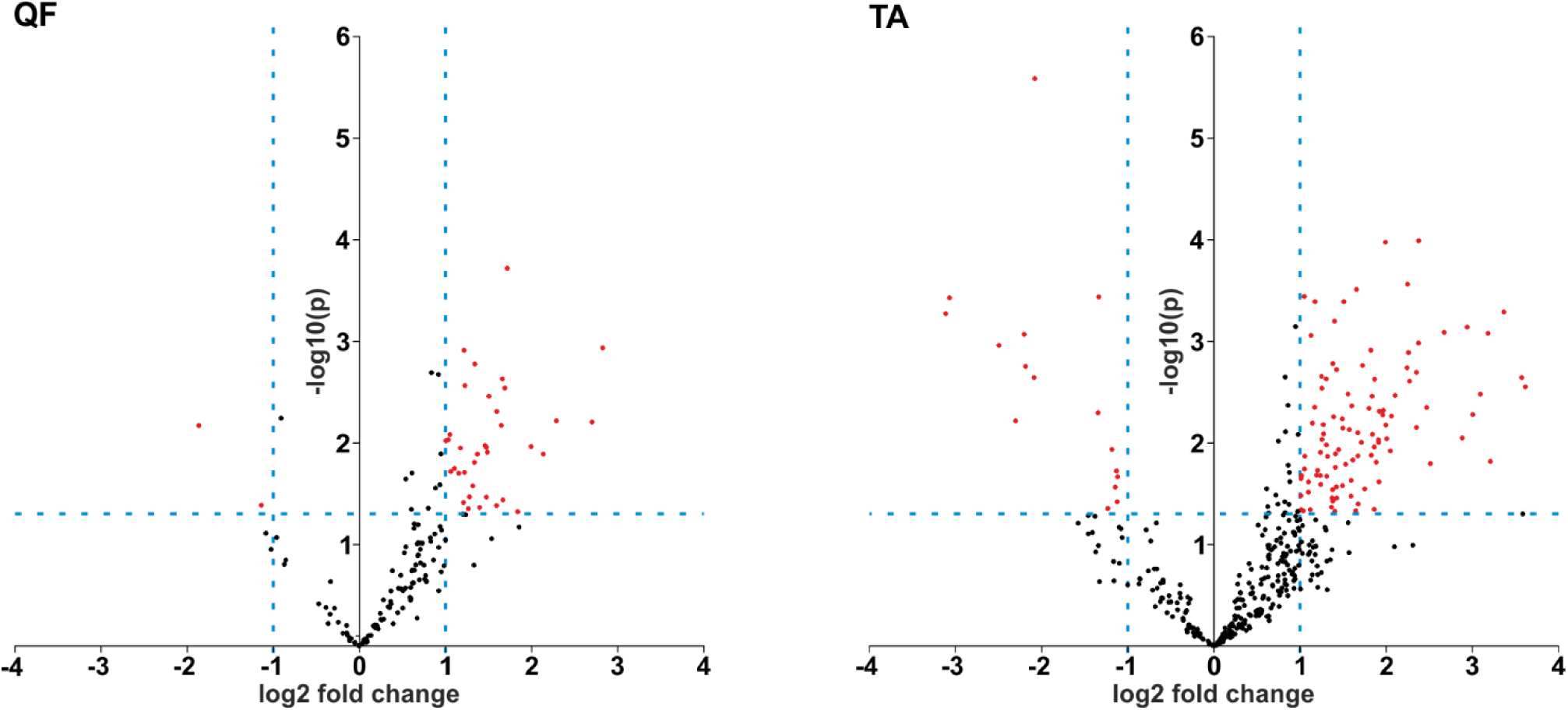
Prevalence of circRNAs with increased levels in DM1. Volcano plots depicting differences in the levels of ‘common’ circRNAs (dots) in DM1 and control samples in QF (left-hand side) and TA (right-hand side). Positive and negative values of log2 fold change indicate increased and decreased circRNAs in DM1. Each red dot represents circRNA fulfilling the following criteria of expression change: p-value<0.05 and log2 fold change ≤-1 or ≥1 (the thresholds indicated by dotted lines).

**Table 4.**
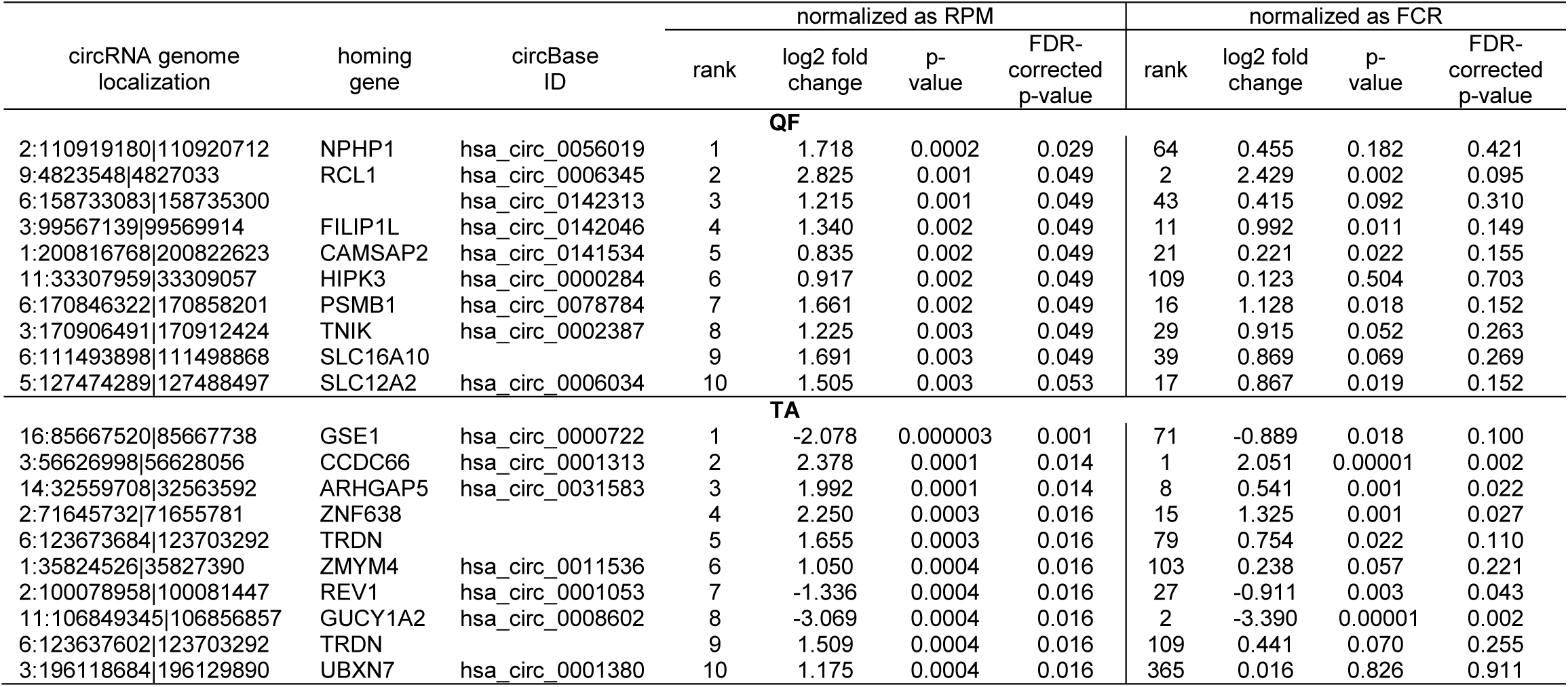
Top ten differentially expressed circRNAs in QF and TA.

The functional association analysis of genes corresponding to circRNAs either increased or decreased in DM1 in TA (67 distinct genes at p<0.01 for differences in RPM, Table S6) showed the strongest association (enrichment) with the following UniProt (UP) keywords: ‘phosphoprotein’ [number of involved genes (n)=46, fold enrichment (FE)=1.7, Benjamini corrected p-value (p_BC_)=0.0005] and ‘alternative splicing’ [n=52, FE=1.5, p_BC_=0.001]. The genes were also associated with the Gene Ontology (GO) cellular component (CC) term ‘nucleoplasm’ [n=24, FE=2.5, p_BC_=0.004]. A similar analysis performed for QF (18 distinct genes) also showed an enrichment of genes associated with alternative splicing and nucleus localization keywords/terms among the top results, but the associations were nonsignificant due to the much smaller number of analyzed genes.

Most of the top differentially expressed circRNAs are deposited in circBase, and the majority of them are encoded by exons of known genes (Tables S3, S4, S5 and S6). The attempted experimental validation of eight randomly selected circRNAs in QF samples confirmed their presence in the studied tissue and the direction of the level change in these circRNAs (in the BP1_DM1_H-sample set; Figure S7).

### Identification of multi-circRNA genes

During the analysis, we noticed that a substantial number of circRNAs were generated from multi-circRNA genes (MCGs), which give rise to more than one circRNA. As shown in Figure 4A, 69% and 78% of circRNAs were generated from MCGs in QF and TA, respectively. Furthermore, 14 MCGs in QF and 59 MCGs in TA (top-MCGs) generated more than ten distinct circRNAs. The top-MCGs from which the highest numbers of circRNAs were generated were *titin* (*TTN*: 44 circRNAs in QF and 86 circRNAs in TA; cumulatively 96 distinct circRNA species), *nebulin* (*NEB*: 41 and 59; cumulatively 66), and *triadin* (*TRDN*: 24 and 37; cumulatively 39). All three genes are strongly related to biological functions and highly expressed in skeletal muscles. Other top-MCGs strongly related to the function of skeletal muscles are *dystrophin* (*DMD*), *myopalladin* (*MYPN*), *myomesin 1* (*MYOM1*), and *myosin IXA* (*MYO9A*). Notably, the abovementioned muscle-related multiexon MCGs were strongly enriched in new (not present in circBase) circRNAs (~95% vs 34%/39% in all ‘validated’ circRNAs in QF/TA samples). This finding may have been observed because skeletal muscle tissues were not comprehensively studied (reported in the circBase) in the context of circRNA discovery.

**Figure 4.**
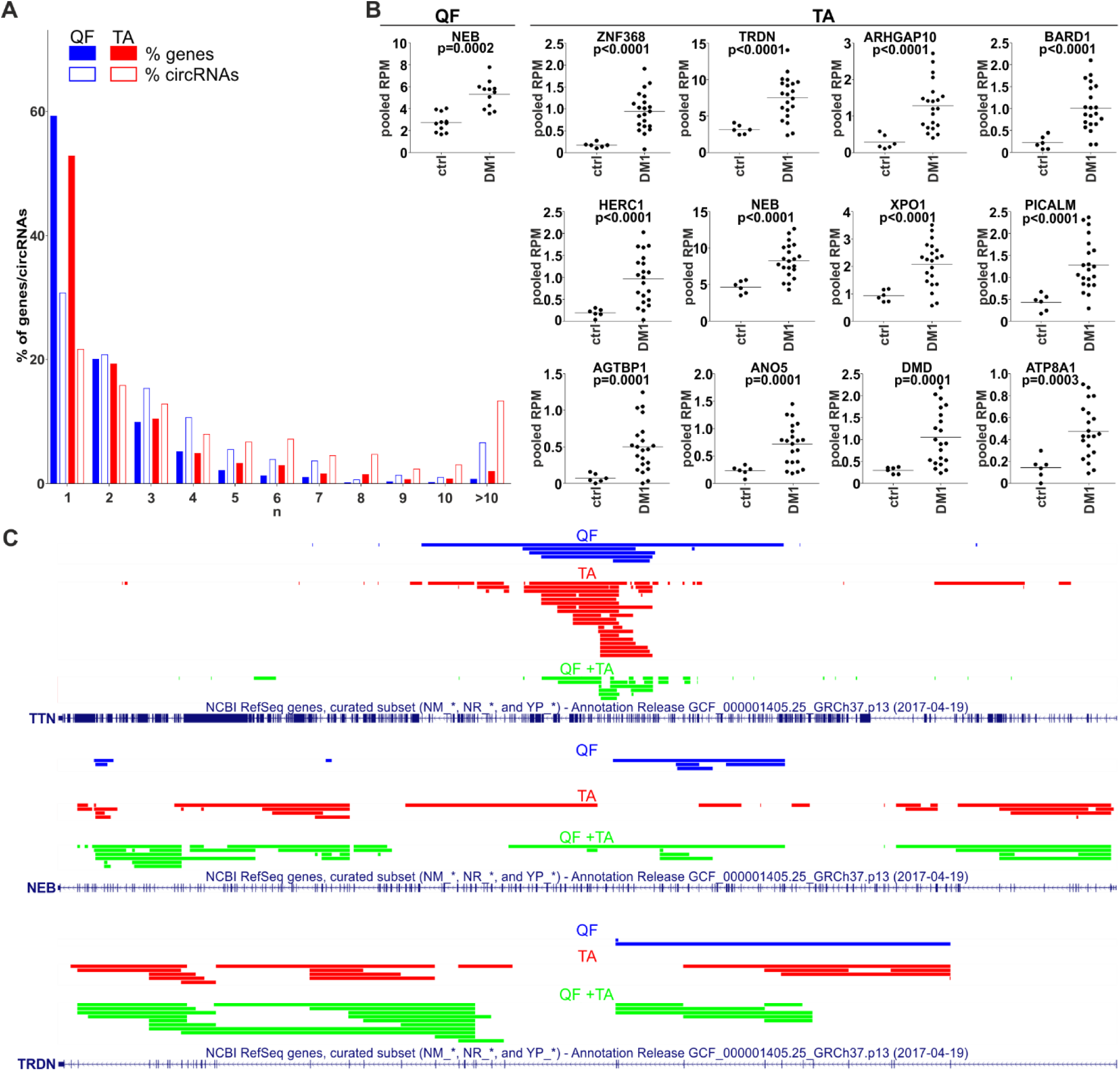
CircRNAs generated from MCGs. **A)** Bar graph showing the percentage of genes that generate a particular number (n) of distinct circRNA species (solid bars) and the percentage of circRNAs generated from these genes (empty bars). Blue and red bars represent QF and TA, respectively. For example, in QF, the genes generating more than ten circRNAs constitute ~1% of all circRNA-generating genes but generate ~7% of all circRNAs. **B)** Dot plots depicting levels (pooled RPMs) of top-MCG-specific circRNA pools most profoundly differentiated between control (ctrl) and DM1 samples in QF and TA. The FDR-corrected p-value is shown above each graph. In each graph, each dot represents pooled circRNA-specific RPM values in the individual sample. **C)** The maps of *TTN*, *NEB* and *TRDN* (RefSeq tracks) with schematic representation of regions (color lines) overlapping exons giving rise to circRNAs (presented with the use of UCSC Genome Browser). Blue, red, and green lines represent circRNAs specific to QF, specific to TA, common to QF and TA, respectively.

The maps of genomic regions giving rise to circRNAs generated from top-MCGs common to QF and TA are shown in Figure 4C and Figure S8. As shown in the figures, the back-splice sites of almost all circRNAs overlapped with the splice sites of canonical exons; therefore, almost all circRNAs may derive from the sequences of canonical exons. Moreover, a substantial fraction of circRNAs were common to QF and TA (green lines, QF+TA), and tissue-specific circRNAs mostly resulted from the higher number of circRNAs detected in TA. Interestingly, in most cases, circRNA-annotated sequences were not randomly distributed and clustered in the center of the gene. The effect was especially visible for circRNAs common to QF and TA. The most profound example of this distribution was *TTN*. The opposite example was *NEB* in which circRNA-annotated sequences were more or less randomly distributed over the entire gene. The observed distributions do not indicate that circRNAs are preferentially generated from exons flanked by long introns (18).

### The level of circRNA pools generated from particular MCGs increases in DM1

Considering circRNAs as competing regulators of linear transcripts, any circRNA generated from a particular gene may affect its linear-transcript-dependent expression. Therefore, in the next step, we compared the cumulative level of circRNAs generated from particular top-MCGs (circRNA pools) in control and DM1 samples. As shown in Table S7 and Table S8, the cumulative RPM value of circRNA pools increased in DM1 samples in 11 out of 14 and 59 out of 59 top-MCGs in QF and TA, respectively. Similar results were also obtained for pooled FCRs (Table S7 and Table S8), as well as for circRNA pools obtained with the other methods of circRNA level normalization (e.g., against the level of housekeeping genes; data not shown). In eight cases (i.e., *GBE1, SMARCC1, BIRC6, SENP6, CHD2, MYBPC1*, *MAP4K3* and *RALGAPA2*), the circRNA pools were increased although none of the individual circRNAs constituting these pools were significantly differentiated. The levels of the most profoundly differentiated circRNA pools (FDR-corrected p-value<0.0005) in QF and TA are shown in Figure 4B.

### CircRNA levels are associated with DM severity

The comparison of the global circRNA level in TA with a phenotypic biomarker of muscle strength (ankle dorsiflexion force) associated with DM1 severity showed a substantial correlation [correlation coefficient (R)=-0.85; p<0.001]. Significant negative correlation with muscle strength (p<0.05; R<-0.434) showed also 117 (out of 403) individual ‘common’ circRNAs and 42 (out of 59) top-MCGs-specific circRNA pools (Figure 5, Table S9).

**Figure 5.**
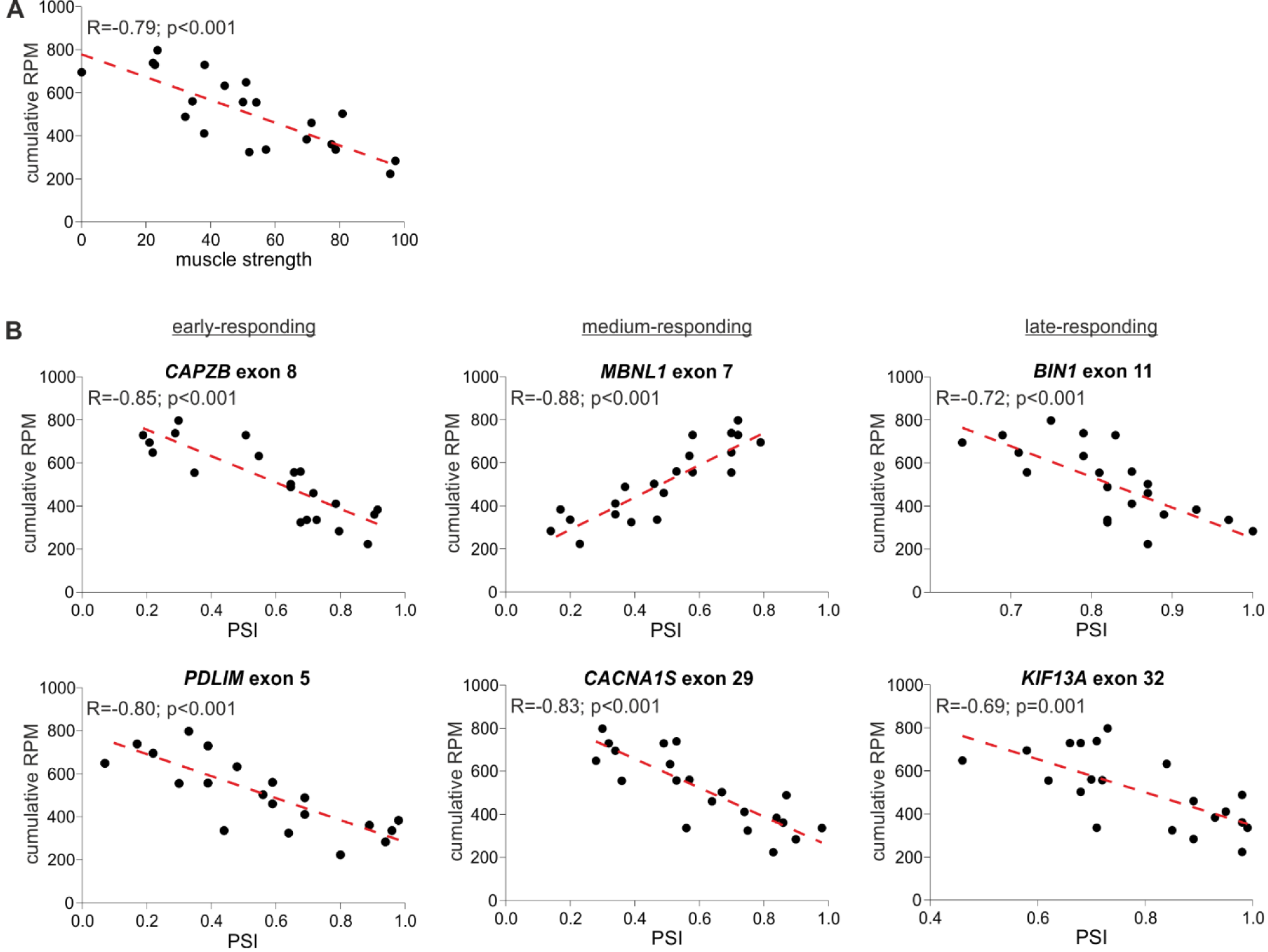
Correlation of global circRNA level with disease severity. **A)** A scatter plot showing correlation of global circRNA levels normalized as RPMs (Y-axis) and muscle strength (X-axis). **B)** Scatter plots showing correlations of global circRNA levels (Y-axis) and PSI values of early-, medium- and late-responding exons alternatively spliced in DM1 (X-axis). For each plot, the R value, p-value and the trendline (red dotted line) are shown, each dot represents an individual TA sample.

In the next step, we compared the circRNA level with the level of early-, medium-, and late-responding alternatively spliced exons, being molecular biomarkers of DM1 severity. As shown in Figure 5 and Table S9, the global circRNA level was significantly correlated with the percent spliced in (PSI) values of all analyzed exons. The strongest correlation showed exon 7 of *MBNL1* (R=0.88; p<0.001), exon 8 of *CAPZB* (R=-0.85; p<0.001) exon 29 of *CACNA1S* (R=-0.83; p<0.001) and exon 22 of ATP2A1 (R=-0.82; p<0.001). Negative correlations were obtained for exons alternatively excluded in DM1. In contrast, exon 7 of *MBNL1* and exon 7 of *NFIX*, both alternatively included in DM1, showed positive correlations. Similar correlations were obtained for a substantial fraction of individual circRNAs, as well as for the top-MCG-specific circRNA pools (Table S9).

## DISCUSSION

Splicing aberrations induced by functional inactivation of MBNL splicing factors constitute a main pathomechanism of DM1. Previous research suggested that in addition to a function in alternative splicing, MBNL proteins participate in the biogenesis of circRNA, bringing circRNA-flanking introns closer together and facilitating back-splicing (circularization) (15). Thus, downregulation of circRNAs would be expected in DM1 (and in DM2) cells in which expanded CUG (CCUG in DM2) repeats attract MBNLs, leading to their sequestration.

To test whether circRNA levels are decreased in DM1 and to verify the role of MBNLs in the biogenesis of circRNA, we analyzed the expression level of up to twenty circRNAs in myoblast cell lines and skeletal muscle samples derived from patients with DM1 and healthy controls. Among the selected circRNAs were those with a relatively high number (n≥10) of potential MBNL-binding motifs in flanking introns, as well as circMBNL1, which is regulated by MBNL1 (37). Additionally, circCDR1as and circHIPK3, the highly expressed and most extensively studied circRNAs, were among the selected circRNAs (17, 23, 24). None of the circRNAs tested in our analysis showed a consistent decrease of level in DM1. There was also no decrease in the levels of circRNAs in muscles from patients with DM2, or in muscles from the transgenic mouse model of DM1. All of the above results question the role of MBNLs as important factors in circRNA biogenesis in muscles. The discrepancy between our study and earlier reports may be because previous analyses were performed in artificial models (artificially generated circRNA genes) in which some of the tested processes (e.g., interaction of MBNLs/Mbl with artificial, usually shorter introns) may take place differently, and the stoichiometry of interacting proteins and RNA particles may be different from those in a natural mammalian tissues. Additionally, the previous experiments were mostly performed with the fly Mbl splicing factor. Potentially, human orthologues may not have the exact same circRNA-generation activity and we cannot exclude the possibility that decreased levels of MBNLs, although they induce aberrations in alternative splicing, are still sufficient for circRNA processing. Furthermore, it is possible that MBNLs play a role in the biogenesis of specific individual circRNAs, which were not tested experimentally in our study. CircGSE1, flanked by multiple MBNL binding motifs and decreased in DM1, may be an example of such a circRNA. MBNL1-dependent biogenesis of circGSE1 may be additionally supported by the fact that opposite to other circRNAs its increased level is associated with lower DM1 severity (Table S9). Another example of circRNA decreased in DM1 and associated with lower DM1 severity is circFGFR1 (Table S9). In contrast to our original hypothesis, the abovementioned experiments showed a trend toward a global increase in circRNA levels in DM1 samples. Although changes in levels of individual circRNAs are small and nonsignificant in most cases, circRNAs with increased levels in DM samples were prevalent in most of our experiments. Additionally, analysis of mouse samples where a higher number of the samples provided a better statistical power to detect smaller changes in circRNA levels showed that two (circCamsap1, and circHipk3) out of the four tested circRNAs were significantly increased in mouse model of DM1.

To check whether the global circRNA level is indeed increased in DM1, we used publicly available RNA-Seq datasets deposited in the DMseq database (http://www.dmseq.org/). The advantage of such data is that they are generated by an independent experimenter blind to the hypotheses tested in particular studies (also in ours). The increased global level of circRNA in DM1 was confirmed in two independent sets of samples, consisting of samples from two different skeletal muscles, QF and TA.

CircRNAs, generated either cotranscriptionally or posttranscriptionally (15, 20, 21), compete with their linear counterparts (mRNAs) for their shared linear precursor (pre-mRNA). However, notably, some circRNAs are the main or exclusive products generated from their precursors (e.g., circCDR1as). The generation of circRNA may be a mechanism of mRNA downregulation (15). Alternatively, disturbances and delays in mRNA maturation may increase the duration of the immature transcript and shift the balance of pre-mRNA processing in favor of circRNA biogenesis (15, 40, 41). In DM1, such disturbances in transcript maturation may be caused by the sequestration of MBNLs and aberrations in splicing. The increased global level of circRNA in DM1 may simply be a side effect of splicing aberrations, or secondary effect of the chronic pathological state of DM1, not dependent on MBNL1 or splicing alterations. Furthermore, as the levels of circRNAs are altered in such disorders as Duchenne muscular dystrophy or dilated cardiomyopathy (42, 43), it may suggest that deregulation of circRNAs is generally associated with a muscle pathological state. However, the possibility that elevated global levels of circRNA or increased levels of an individual circRNA may also play a role in DM1 pathogenesis cannot be excluded. It is supported by the fact, that the global circRNA level, as well as the levels of substantial fractions of MCG-specific circRNA pools and individual circRNAs were negatively correlated with molecular and clinical biomarkers of DM1 severity. Additionally, recent results by Voellenkle et al., that have been made public as a preprint (https://doi.org/10.1101/452391) during the final steps of preparation of our manuscript, showed that the levels of four out of nine tested circRNAs were significantly increased in DM1 patients and correlated with muscle weakness.

By using generated circRNA datasets, we also performed analyses of individual circRNAs and MCG-specific circRNA pools. The analyses led to the identification of many circRNAs and circRNA pools that were significantly differentiated between DM1 and control samples. In both types of analyses and in both analyzed tissues, there was a substantial excess of circRNAs or circRNA pools in DM1. This finding is consistent with the observation of the global increase in circRNA levels in DM1 samples. Although many of the changes in circRNA and circRNA pools reached statistical significance (p<0.05, even after FDR-correction), whether the differentiated circRNAs/circRNA pools are specific and biologically relevant to DM1 or result from a global increase in circRNA levels in DM1 cannot be established. One hint as to the role of circRNAs in DM1 may be found in the functional association analysis, which showed that terms related to alternative splicing and nuclear localization were among the strongest associations of genes giving rise to differentiated circRNAs. Other links between aberrations in circRNA levels and DM1 pathogenesis come from the observed associations between circRNA levels and muscle weakness, as well as between circRNA levels and abnormalities of alternative splicing of well-known DM biomarkers. Additionally, the transcripts of at least 10 (*DMD, KIF1B, MYBPC1, NEB, NCOR2, PICALM, RERE, SMARCC1, UBAP2* and *USP25*) out of 63 identified top-MCGs were previously shown to be aberrantly spliced in DM1 (44, 45). Nonetheless, the changes in individual circRNAs require further experimental validation. Moreover, notably, the power of this analysis is limited due to the depth of coverage (adjusted for mRNA analysis) that does not allow reliable estimation of low-level circRNAs.

Interestingly, among the top-MCGs, there are genes highly expressed and strongly associated with the biological function of skeletal muscles [e.g., *TTN* (total number of circRNAs generated in both QF and TA, n=96), *NEB* (n=66), *TRDN* (n=39), *DMD* (n=33), *MYPN* (n=22), *MYOM1* (n=18), *or MYO9A* (n=14)]. All of these genes are large multiexon genes, including *DMD* (2.1 Mbp, up to 81 exons), the largest human gene, and *TTN* (0.3 Mbp, up to 362 exons), which has the highest number of exons (Figure 4C and Figure S8). A large number of exons increases the number of potential splicing donor/acceptor pairs, which may facilitate the generation of different circRNAs. Alternatively, the higher number of circRNAs generated from multiexon genes may also result from higher chances/numbers of aberrations occurring during processing of their transcripts.

In conclusion, our results indicate that MBNL deficiency does not cause the expected decrease in circRNA levels in DM1 cells and tissues. In contrast, the global level of circRNAs is elevated in DM1. However, the role of the increased level of circRNAs in the pathogenesis of DM1 is unknown and requires further investigation.

## MATERIALS AND METHODS

### cDNA samples

Seven cDNA sample sets (Table 5 and described below) were used in this study. These sets included samples from myoblast cell lines (CL) derived from human skeletal muscles, muscle biopsy (BP) samples from DM1 and DM2 patients and corresponding healthy controls, and samples from the *HSA*^LR^ transgenic mouse model of DM1 (MM). For the purpose of cDNA generation, total RNA was extracted using the standard protocol, as previously described (46). Reverse transcription was performed according to the manufacturers’ recommendations with the use of reverse transcriptases (RTs) either with (H+ sample sets) or without (H-sample sets) RNase H activity. All reverse transcription reactions were performed with the use of random hexamers. The particular RTs used in the analyzed sample sets are indicated below. The DM1-specific splicing aberrations in the muscle sample sets used in this study were evaluated before (47) and are shown (BP3_DM1_H- and BP2_DM2_H-) in Figure S1. The splicing aberrations in DM1 samples deposited in the DMseq database and analyzed in this study (see subchapter Analysis of NGS data) were also recently demonstrated (48).

**Table 5.** Characteristics of sample sets used in the study.

|  | Sample set ID | Subset | # of samples | RNase H activity |
| --- | --- | --- | --- | --- |
| CL | CL1_DM1_H- | control | 3 | H- |
|  |  | DM1 | 3 |  |
|  | CL2_DM1_H+ | control | 2 | H+ |
|  |  | DM1 | 3 |  |
| BP | BP1_DM1_H- | control | 2 | H- |
|  |  | DM1 | 8 |  |
|  | BP2_DM1_H+ | control | 6 | H+ |
|  |  | DM1 | 5 |  |
|  | BP3_DM1_H- | control | 4 | H- |
|  |  | DM1 | 2 |  |
|  | BP4_DM2_H- | control | 4 | H- |
|  |  | DM2 | 9 |  |
| MM | MM_DM1_H- | control, *FVB* | 10 | H- |
|  |  | DM1 model, *HSA<sup>LR</sup>* | 10 |  |

CL–human cell lines; BP–human muscle biopsies; MM–mouse muscles

The sample sets: (i) CL1_DM1_H-(generated with SuperScript III RT, Invitrogen, Carlsbad, CA, USA) consisted of 3 DM1 samples extracted from DM1 myoblast cell lines (9886, >200 CTG repeats; 10010, >200 CTGs; and 10011, >350 CTGs) and 3 sex- and age-matched control samples extracted from non-DM myoblast cell lines (9648, 10104, 10701) as described in (49); (ii) CL2_DM1_H+ (iScript RT, Bio-Rad, Hercules, CA, USA) consisted of 3 DM1 samples (from cell lines 9886, 10010 and 10011) and 2 control samples (from cell lines 10104 and 10701); (iii) BP1_DM1_H-(SuperScript III RT, Invitrogen) consisted of 8 DM1 and 2 control samples. All samples were derived from the QF muscles of either patients with DM1 or non-DM individuals. Samples differed in the number of CTG repeats in the *DMPK* 3’-UTR region, ranging from 50 to 1456 repeats [for details please see (47)]; (iv) BP2_DM1_H+ (iScript RT, Bio-Rad) consisted of 5 DM1 and 6 control samples; (v) BP3_DM1_H-(GoScript RT, Promega) consisted of 2 DM1 and 4 control samples, derived from skeletal muscle tissue; (vi) BP4_DM2_H-(GoScript RT, Promega) consisted of 9 DM2 and 4 control samples, derived from skeletal muscle tissue. Importantly, control samples were the same as those in the BP3_DM1_H-sample set; (vii) MM_DM1_H-(SuperScript III RT, Invitrogen) consisted of 10 DM1-model and 10 control samples of the *HSA*^LR^ transgenic mouse model of DM1 and control background *FVB* mice, respectively. RNA was extracted from gastrocnemius muscle (29).

The samples, experimental protocols, and methods reported in this study were carried out in accordance with the approval of the local ethics committees: NRESCommittee.EastMidlands-Nottingham2 and the University of Rochester Research Subjects Review Board. Informed consent was obtained from all subjects.

### Selection of circRNAs for experimental analyses

Twenty circRNAs (Table 1) whose levels were experimentally evaluated in our study were selected from previously detected (16-18, 22, 40) circRNAs deposited in circBase (December 2016) [(35); http://www.circbase.org/]. We considered only circRNAs validated by at least twenty next-generation sequencing (NGS) reads in at least two of the abovementioned studies. Fourteen circRNAs were selected based on the relatively high level in different types of cells/tissues and a relatively high [≥10% in (18)] proportion compared to that of their linear counterparts (mRNA). Four circRNAs were selected based on a high number (n≥10) of potential MBNL-binding sites [YGCY motifs; (36)] in adjacent (300 nt upstream and 300 nt downstream) sequences of their flanking introns. Two additional circRNAs selected for analysis were circCDR1as and circMBNL1. Additionally, eight circRNAs were experimentally analyzed for the purpose of validation of the most differentiated circRNAs identified based on RNA-Seq data analysis of control and DM1 QF samples (see below).

### PCR assays design and validation

For the experimental analysis of selected circRNAs, we designed PCR assays that allowed the amplification and parallel analysis of circRNAs and their linear counterparts. Each assay consisted of one primer common to the circular and linear transcript and two primers specific for either circular or linear transcript. The only exceptions were assays designed for circCDR1as (circRNA generated from a single-exon transcript) and circMBNL1, which consisted of four primers (two for the circular transcript and two for the linear transcript). Primer sequences are shown in Table S1.

The PCR products of the designed assays were validated by analysis in agarose gel electrophoresis (the length of each product was as expected). Briefly, PCR was performed in a 10-μl reaction composed of 0.3 μl of a 10 μM dilution of forward and reverse primers (0.6 μl in total; primers were synthesized by Sigma-Aldrich, Saint Louis, MO, USA), 0.125 μl dNTP mix (concentration of each nucleotide was 10 mM) (Promega), 0.05 μl GoTaq DNA Polymerase (concentration 5 u/μl) (Promega), 2 μl 5X colorless GoTaq reaction buffer (containing 7.5 mM MgCl_2_) (Promega), 6.225 μl deionized water, and 1 μl cDNA template. The following cycling conditions were used: 2 min at 95°C, followed by 35 cycles at 95°C for 20 sec, 58-60°C (different for individual assays) for 20 sec, and 72°C for 20 sec, followed by 5 min at 72°C. The obtained PCR products were visualized on a standard 1.5% agarose gel. Additionally, the specificity of each product was confirmed by Sanger sequencing performed on an ABI Prism 3130 genetic analyzer (Applied Biosystems, Carlsbad, CA, USA) according to the manufacturer’s general recommendations.

### Droplet digital PCR (ddPCR)

The level of circRNAs was analyzed with the use of the ddPCR technique (32, 33) developed by Bio-Rad. Analyses were performed according to the manufacturer’s general recommendations. Briefly, reactions were carried out in a total volume of 20 μL, containing 10 μL 2X EvaGreen Supermix (Bio-Rad), 1 μL 4 μM forward primer, 1 μL 4 μM reverse primer and different amounts of cDNA template, determined on the basis of optimization reactions performed for each analyzed gene/transcript. A QX200 ddPCR droplet generator (Bio-Rad) was used to divide the reaction mixture into up to 20,000 droplets. The initial dilution of the cDNA samples ensured that most of the generated droplets contained zero or one template molecule. The thermal parameters of the PCR were as follows: 5 min at 95°C, followed by 40 cycles of 30 sec at 95°C, 30 sec at annealing temperature (optimized for each gene) and 45 sec at 72°C, followed by 2 min at 72°C, 5 min at 4°C, enzyme inactivation at 90°C for 5 min and holding at 12°C. The amplified products were analyzed using a QX200 droplet reader (Bio-Rad). The exact number of cDNA particles (representing particular transcripts) was calculated based on the number of positive (containing template cDNA molecules) and negative (without template cDNA molecules) droplets using QuantaSoft (Bio-Rad) version 1.7.4.019 software, which utilizes Poisson distribution statistics.

In the analyses, we took the factor of the aforementioned cDNA dilution into account. Importantly, in our analysis, we used the following exclusion criteria: (i) from the analysis of the level of a particular circRNA, we excluded samples with less than ten positive droplets corresponding to the linear counterpart of this circRNA; (ii) in the individual sample set, we did not consider the analysis of a particular circRNA if more than half of the samples included in this sample set were excluded from the analysis in step (i). Additionally, due to the limited amount of RNA samples, not all of the originally selected circRNAs were tested in the BP3_DM1_H- and BP4_DM2_H-sample sets.

For each analyzed circRNA, their levels in particular samples were calculated as a fraction of circular particles (FCP) constituted by the amount of circRNA particles (C) in a total number of particles [circRNAs (C) and their linear counterparts (L)] generated from a particular gene:

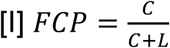

The only exception was circCDR1as for which both linear and circular transcripts are generated from the same single exon (PCR primers designed for analysis of linear transcripts are also specific to cDNA generated from circular transcripts). Thus, the equation in this case is as follows:

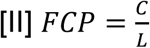

Additionally, the levels of circRNAs and their linear counterparts were normalized against the levels of housekeeping genes (i.e., *ACTB* and *GAPDH*).

### Analysis of NGS data

For the purpose of global analysis of circRNA expression, we used the RNA-Seq data [GEO (GSE86356)] deposited in the DMseq database (48) (http://www.dmseq.org/). From the data sets of 126 samples derived from different muscle tissues, we chose the data sets of muscles represented by the highest number of samples, i.e., QF and TA. To avoid potential technical variations for analysis, we selected only samples for which sequencing data were generated with uniform procedures. For each sample, paired-end sequencing libraries were prepared from rRNA-depleted total RNA. Reverse transcription was performed using random primers, followed by second strand cDNA synthesis, end repair, adenylation, and ligation of adapters. Sequencing was performed using an Illumina HiSeq 2000 system (Illumina, San Diego, CA, USA), followed by processing with standard HiSeq 2000 software. Reads were mapped to the human genome (GRCh37/hg19) using Hisat2 (50). For the analysis, we selected data sets for 23 QF samples (11 control samples and 12 DM1 samples) and 27 TA samples (6 control samples and 21 DM1 samples). The GSM accession numbers of selected samples are as follows: QF–GSM2309550, GSM2309551, GSM2309552, GSM2309553, GSM2309554, GSM2309555, GSM2309556, GSM2309557, GSM2309558, GSM2309559, GSM2309560, GSM2309565, GSM2309566, GSM2309567, GSM2309568, GSM2309570, GSM2309571, GSM2309572, GSM2309573, GSM2309574, GSM2309575, GSM2309576, GSM2309577; TA–GSM2309586, GSM2309587, GSM2309592, GSM2309593, GSM2309594, GSM2309595, GSM2309599, GSM2309601, GSM2309603, GSM2309604, GSM2309605, GSM2309606, GSM2309611, GSM2309612, GSM2309613, GSM2309616, GSM2309617, GSM2309619, GSM2309621, GSM2309622, GSM2309624, GSM2309629, GSM2309632, GSM2309635, GSM2309639, GSM2309640, and GSM2309641. The average number of mappable reads in selected samples was ~29 million (ranging from ~18 to ~97 million reads; median ~26 million reads) and constituted 92% of the total library size on average. The length of reads was 60 nt. The detection and quantification of circRNAs and their linear mRNA counterparts in the selected samples was performed with CIRI2 (38). The normalized level of circRNAs was calculated either as a number of circRNA-specific reads per million mappable reads (RPM) or as a fraction of circRNA-specific reads in a total number of circRNA-specific and corresponding linear reads (FCR). Note that FCR corresponds to FCP calculated based on the number of circular and linear RNA particles. The level of circRNAs was also normalized against the number of reads specific to individual housekeeping genes (e.g., *ACTB* or *GAPDH*).

### Statistical information

All statistical analyses were performed using Statistica (StatSoft, Tulsa, OK, USA) or Prism v. 5.0 (GraphPad, San Diego, CA, USA). All p-values were provided for two-sided tests. If necessary, the false discovery ratio (FDR) was calculated according to the Benjamini-Hochberg procedure (http://www.biostathandbook.com/multiplecomparisons.html). All human genome positions indicated in this report refer to the February 2009 (GRCh37/hg19) human reference sequence. The functional association analysis of the genes corresponding to circRNAs was performed with the use of DAVID Bioinformatics Resources (51, 52). Correlations of circRNA levels with DM1 severity were performed for TA samples with the use of phenotypic (ankle dorsiflexion force) and splicing alteration data deposited in the DMseq database.

