## Supplemental Figures and Supplemental Tables S1-S2 for "Global increase in circRNA levels in myotonic dystrophy"

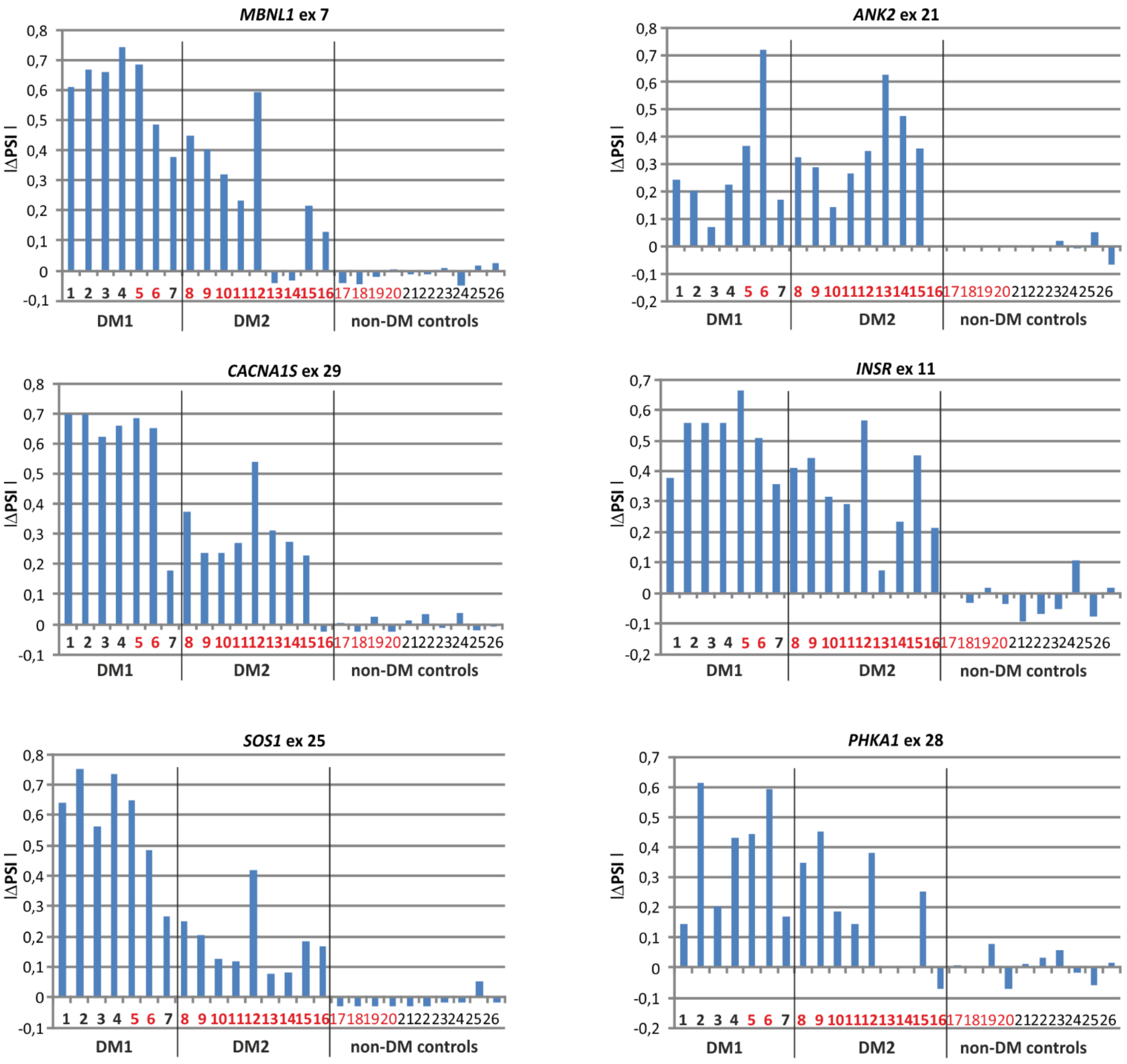


**Figure S1. A)** Splicing changes of alternative exons of six genes in DM1 (samples 1-7) and DM2 (samples 8-16) compared to non-DM muscle samples. For each DM sample the deltaPSI was calculated based on MLPA-based splicing assays described before (47). Samples included in BP3_DM1_H- and BP4_DM2_H- sample sets are indicated in red.

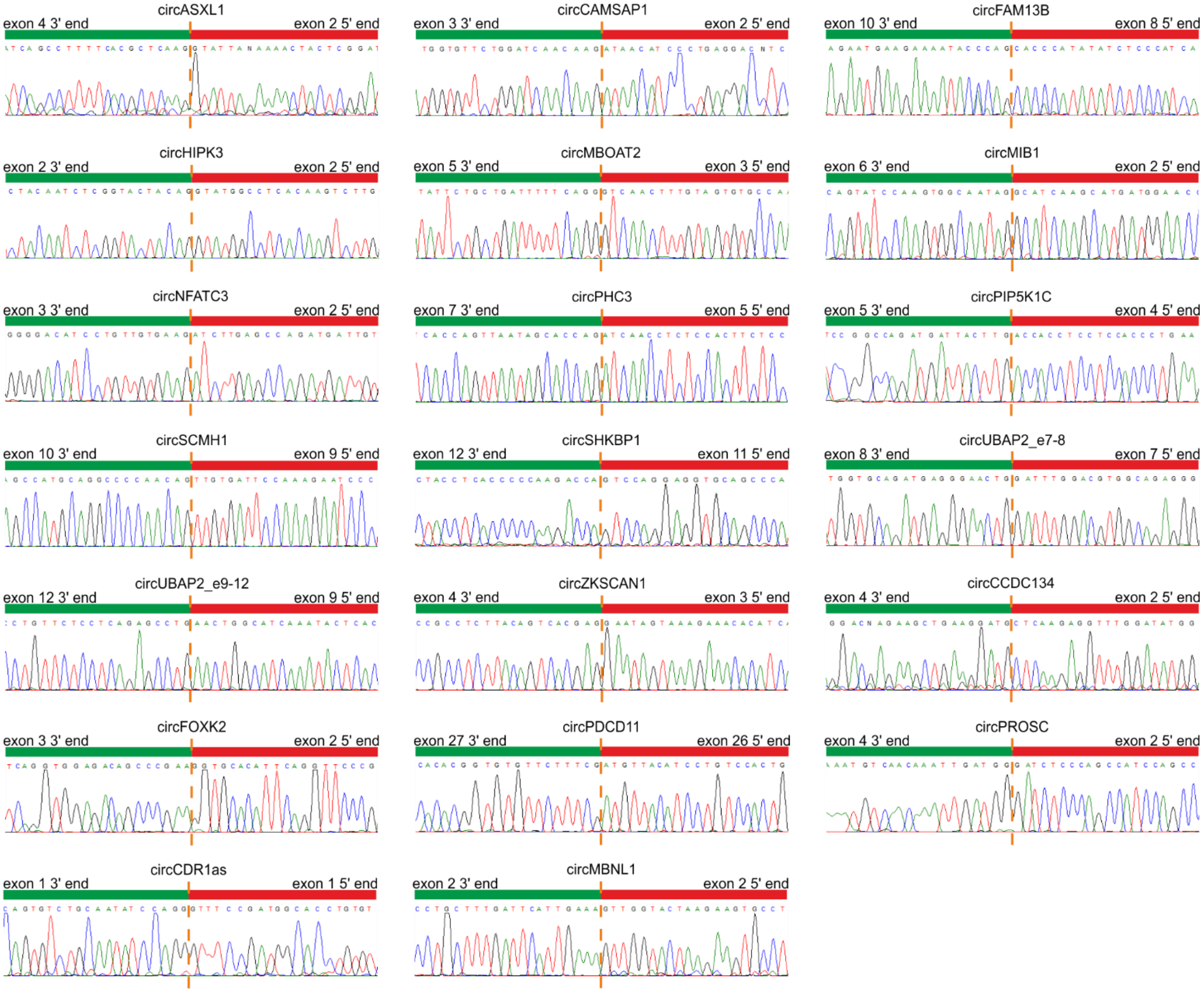


**Figure S2.** Results of Sanger sequencing of the predicted back-splice sites of the selected circRNAs.

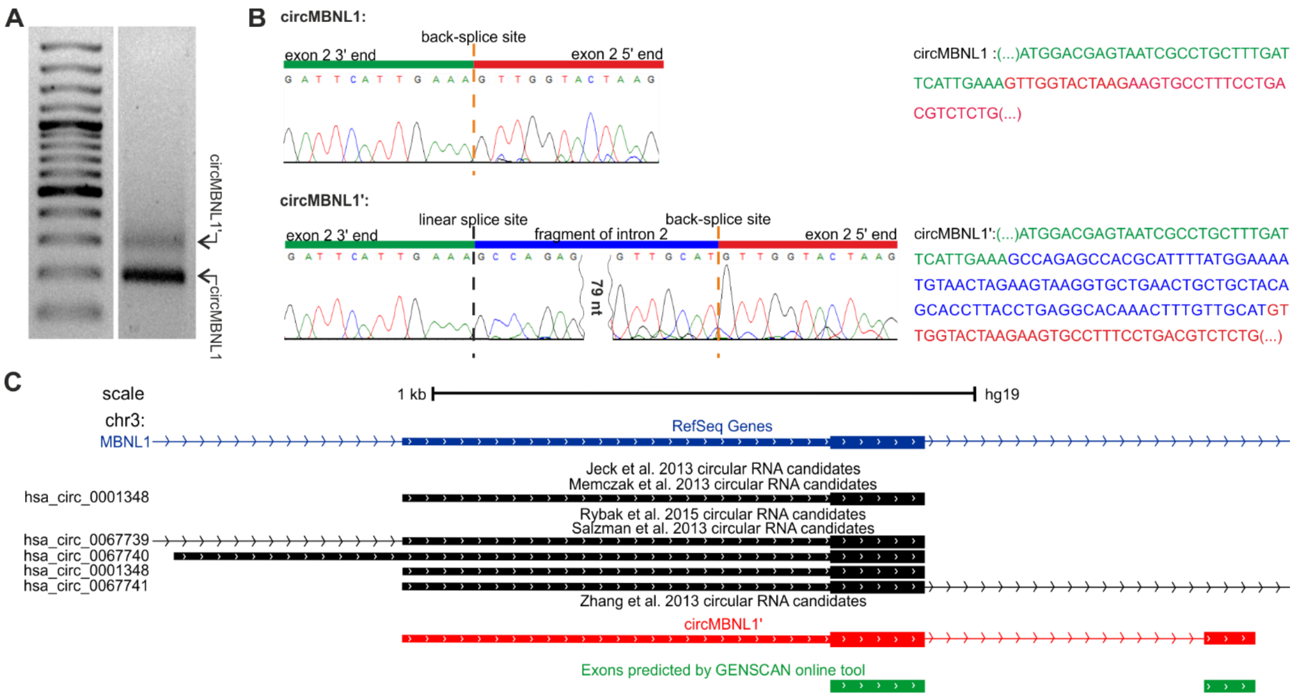


**Figure S3.** Identification of a new circRNA derived from the *MBNL1* gene. **A)** Agarose gel electrophoresis of the product of PCR performed with the use of circMBNL1-specific primers (right-hand side track). Lower and upper bands represent known circMBNL1 (hsa_circ_0001348) and the newly identified circMBNL1', respectively. The size of the upper band does not correspond to a potential concatemer. The left-hand side track is the GeneRuler 1 kb DNA Ladder (Thermo Fisher Scientific). **B)** The results of the Sanger sequencing of PCR products corresponding to upper and lower bands. Sequencing of the lower band confirmed the expected circMBNL1 back-splice site. However, sequencing of the upper band revealed an additional 93-nt long fragment of the downstream intron. The sequences of both circRNAs are shown to the right. **C)** The map showing the localization of MBNL1 circRNAs. The blue (RefSeq) track shows a fragment of *MBNL1* overlapping exon 2. Black tracks indicate known circRNAs deposited in the circBase. The red track indicates the localization of newly identified circMBNL1'. The green track depicts exons predicted with high confidence by the GENSCAN online tool.

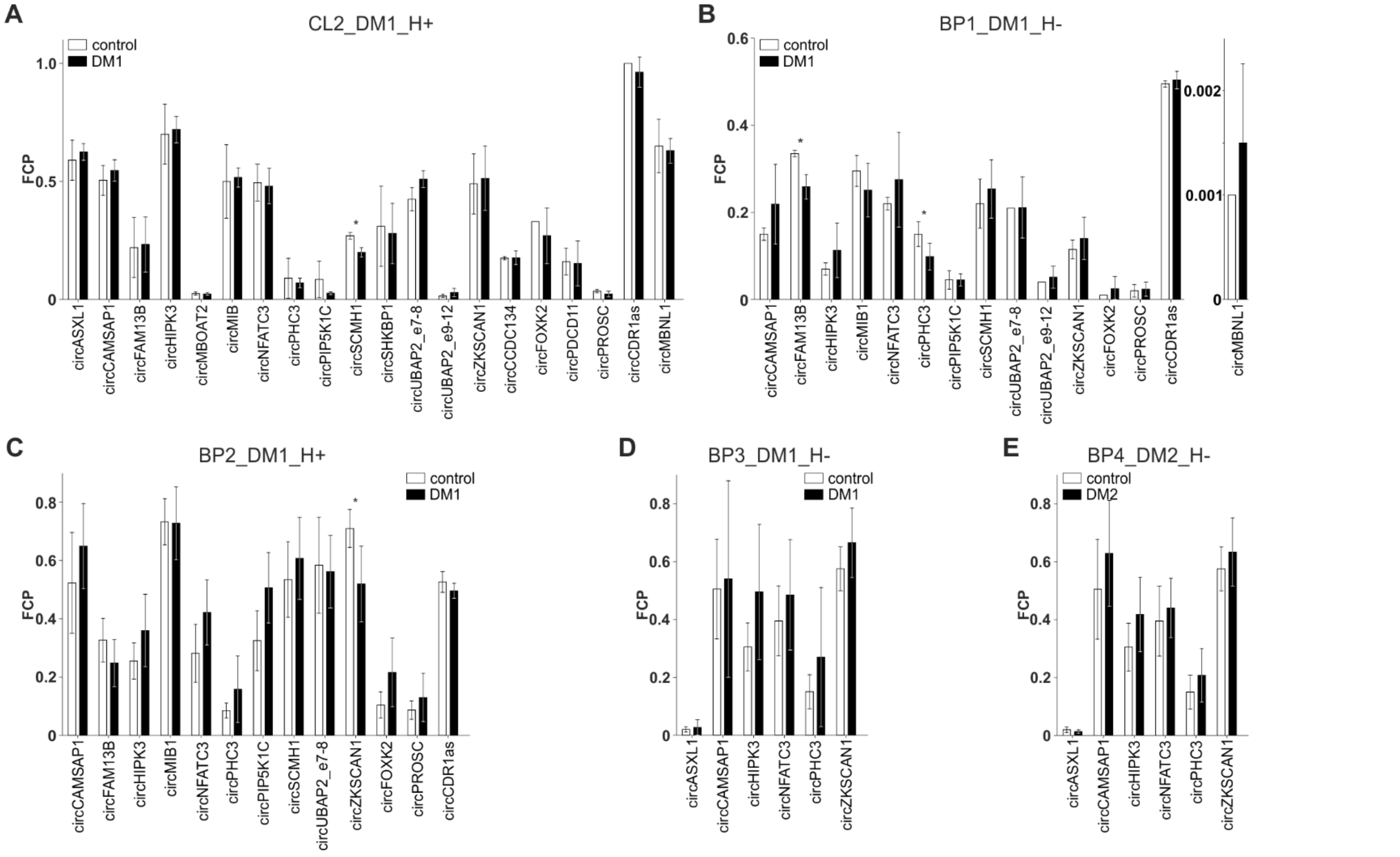


**Figure S4**. Bar graphs showing the results of circRNA expression analysis performed with the use of different sample sets. **A)** CL2_DM1_H+, **B)** BP1_DM1_H-, **C)** BP2_DM1_H+, **D)** BP3_DM1_H-, **E)** BP4_DM2_H-. The scheme of the bar graphs is similar to that used in Figure 1E.

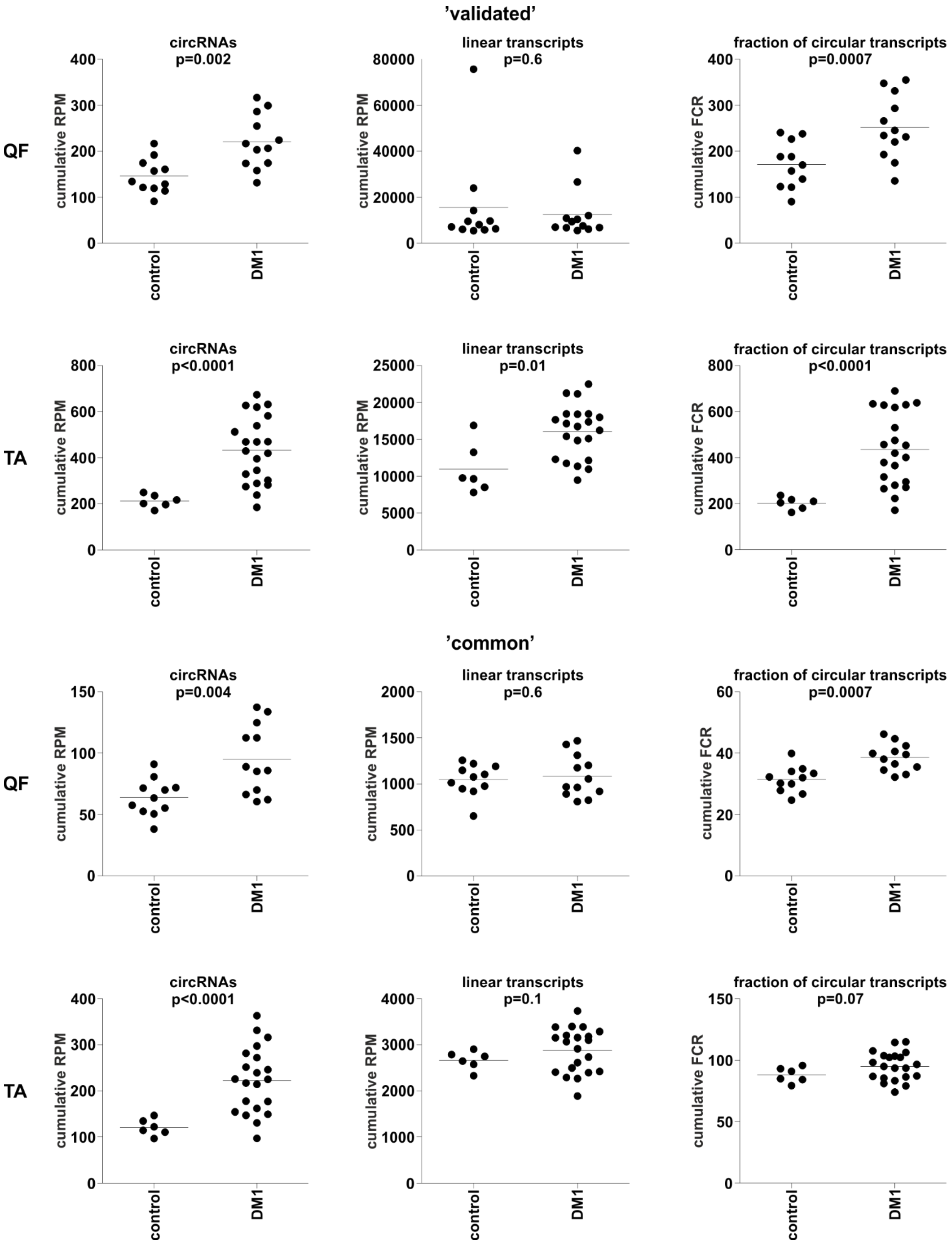


**Figure S5.** Dot plots depicting the cumulative level of ’validated’ (upper panels) and ‘common’ (lower panels) circRNAs in control and DM1 samples of QF and TA. The scheme of the panels is the same as that used in Figure 2.

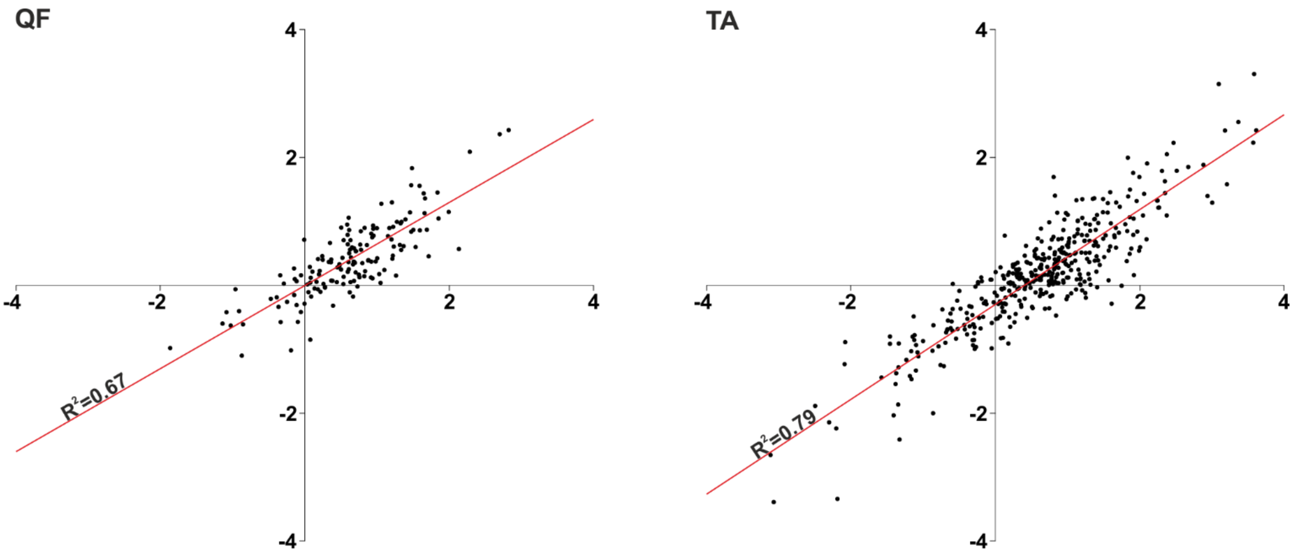


**Figure S6.** Scatter plots showing correlations of circRNA levels normalized as RPMs (axis X) and FCRs (axis Y). For both QF (on the left-hand side) and TA (right-hand side) samples, the $R^2$ values are shown above the trendline (red line). Each dot represents an individual circRNA.

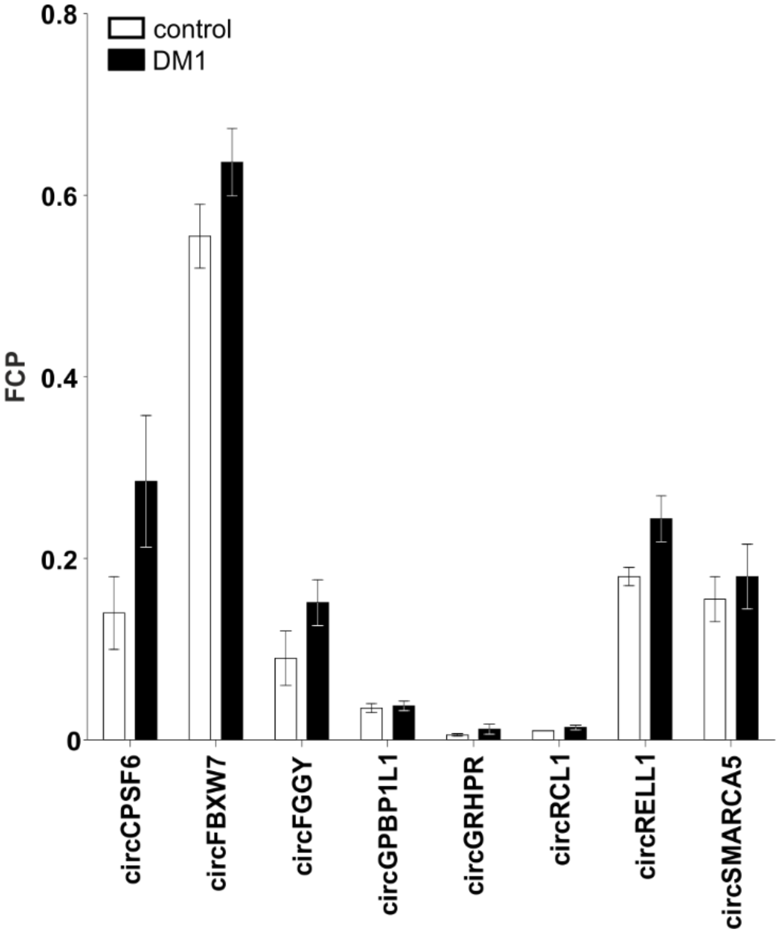


**Figure S7.** Bar graph showing results of ddPCR analysis of the level of eight circRNAs randomly selected from ‘common’ circRNAs of QA muscle. The analysis was performed using the muscle biopsy (BP1_DM1_H-) sample set. The scheme of the bar graph is similar to that used in Figure 1E. The results confirmed the direction of expression changes in circRNAs upregulated in DM1 in QA.

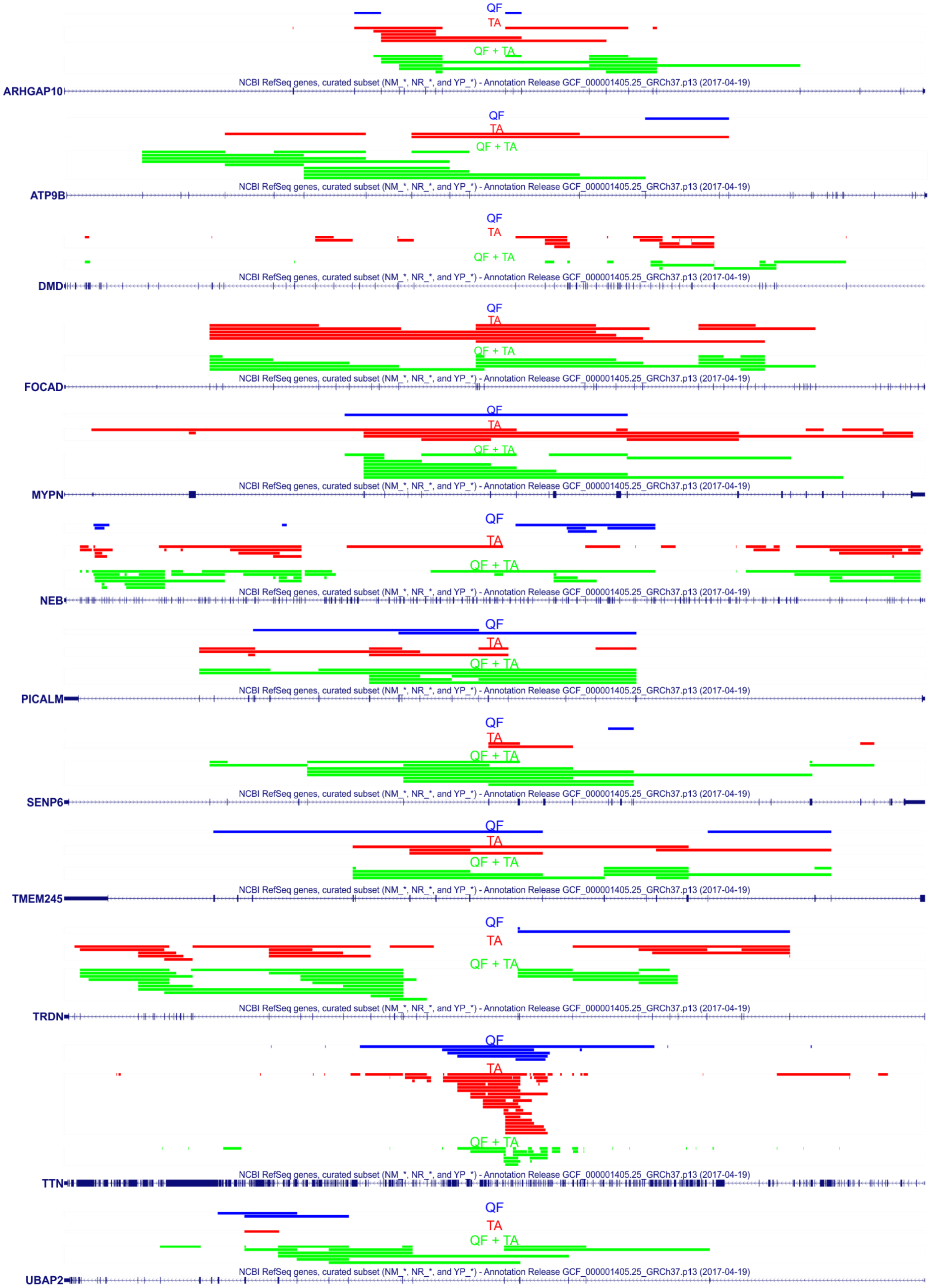


**Figure S8.** The maps of genomic regions of the top-MCGs in which both QF and TA consequently generate more than ten distinct circRNA species. The colors of the tracks are shown in Figure 4C.

**Supplementary Table S1. PCR primers used in the study.**

| gene | circular/linear transcript | primer forward sequence | primer reverse sequence |
|---|---|---|---|
| | | **primers designed for analyses of human circRNAs** | |
| **ASXL1** | cicrular<br>linear | TGCCTCAATGCTATGCTACA | TTTAGTCCTTCTGCCTCTATGA<br>CGACTCTGAGATTCTGTAGAAC |
| **CAMSAP1** | cicrular<br>linear | CCTACACTGTGGAGATGATC | TGATAACAGGCGGCTTAATG<br>CTCATCAAATCCTCCAACAA |
| **CCDC134** | cicrular<br>linear | ATCCTTGATGTCATGCTCAAG | AAAGCAGGACAAAGAGGAAG<br>AAAGTAATAGTGCACAATCCTCG |
| **CDR1as** | cicrular<br>linear | CCATCAACTGGCTCAATATC<br>GCCAATATATGTCTTCCTGAAG | AGGATTATCTGGAAGACCTTGA<br>AAGACCCAAATTATCCGGAA |
| **FAM13B** | cicrular<br>linear | GCAGAAAGTATTCATTGTGATG<br>AAGCTGGAACATATTGAAGAACTT | GTAGGATGCTGATGGGAGAT |
| **FBXW7** | cicrular<br>linear | TCTGACCCAGTAACTCCACT | TCAGTCTGATGATAGTAGCAGAGA<br>AAGGAGGAAGGGAACCAA |
| **FGGY** | cicrular<br>linear | TCAATGAGACCAAGCACAGT | TCAAACCCAAGTCCTCGAAT<br>TTGCCTTCCACGATAAGAAG |
| **FOXK2** | cicrular<br>linear | TGCCATCTGACCTCAATTTAA | TGCTTCTCTCTCTTCTCGCT<br>GCAGTCCTGTAGTAGGGATAAT |
| **GPBP1L1** | cicrular<br>linear | AATGGGAACAAATTGTCATCC | AATAGGTCTGCATGGCTGAT<br>GGTAACAGAGATTGGACTGG |
| **GRHPR** | cicrular<br>linear | AGATGTCCTGACAGATACCAC | ACACCTCGCTCTAGCTCCTT<br>TGTGTACAGAAATCTCTGGACA |
| **HIPK3** | cicrular<br>linear | TATGTTGGTGGATCCTGTTC | ATTCACATAGGTCCGTGGATA<br>AATAATTCTGCAATCACACATCC |
| **MBNL1** | cicrular<br>linear | GGAAGTATGTAGAGAGTTCCAG<br>CAAGTTGAAAATGGACGAGTA | AAGCAGAGACTGCTCTAGAAG<br>ATGTTCTTCTGCTGAATCAAGTT |
| **MBOAT2** | cicrular<br>linear | TCTGGGATACCTCACAGTGT<br>CATCGACCAGGTCAACTTT | GGGTAGCAACTACATGTCTTATAA |
| **MIB1** | cicrular<br>linear | GACTGATGGAATGTTTGAGAC | AACACACTGTGCACAAATCAT<br>ATCTCCACTTCGGACAATGT |
| **NFATC3** | cicrular<br>linear | ATGAAACTGAAGGTAGCCGA | TGTGGTAAGCAAAGTGGTGT<br>TGGCAATTATTATCTCTTGGCTT |
| **PDCD11** | cicrular<br>linear | ATCCAGAGATTAACTCCATCCA | CTACTTTGCTTTTCGTCTCC<br>GAGGTGTTTGTTATAAAGGGC |
| **PHC3** | cicrular<br>linear | TAATAAGAAAGGAGAGAGCCCA<br>AGCAGTCAAGTATGTCCCAA | GAGATACATTTGAGCTTGGC |
| **PIP5K1C** | cicrular<br>linear | TCAAGACCTATGCACCTGTC | ACGTAGAAGTCCTGCATGAG<br>GCAGTACAGCCCATAGAACT |
| **PROSC** | cicrular<br>linear | TTCTGTCTTTGTGTCCTGAGAT | ATCACCATGTCTGCAGGTTT<br>TCTTCTCCGCTGGTGTTAAT |
| **RCL1** | cicrular<br>linear | CTGGAGAGTCTTCTTTGCTT | TTATTTCAATTCGAGAACCATTC<br>TCAGTTCAAATGATTCACCATC |
| **RELL1** | cicrular<br>linear | AGGCTGTTATCTGCTACCATC | AAAGTGGAGCACAAGTCAAAC<br>GAGCAAGATATCGAAGAGGAA |
| **SCMH1** | cicrular<br>linear | AACAACCAGCACTCCTGAAC | ATGGGAAATTAGGGTCTTGG<br>AGTTGCTGGACCTTCTTCTTAT |
| **SHKBP1** | cicrular<br>linear | ATGAAAGACAACGACCTCCT | TAAATGGAGCCGTTGTTGCA<br>TTCTCCGACAGCATGATCTT |
| **UBAP2_ e7-8** | cicrular<br>linear | ACATTTAATCCTGCAGACTATTCA<br>AAAGCGAGAAAGAATCGAGTC | ATGCCTTGGGTTGAGAACCT |
| **UBAP2_ e9-12** | cicrular<br>linear | ATTGATCTGGTAGCCTTGCT<br>CATTTAATCCTGCAGACTATTCA | TCCTGAGCTATGTTGTGAGTATT |
| **ZKSCAN1** | cicrular<br>linear | TCTGGATCCAGTTCAGGAGT | GTCTGTGCTCACCTTTATGTC<br>TTGTCCCTACTGAGATTCCTC |
| | | **primers designed for analyses of mouse circRNAs** | |
| **Camsap1** | cicrular<br>linear | CCTACACTGTGGAGATGATC | TGATAACGGGTGGCTTAATG<br>GATTTACCTTATTGATCCAGAATACC |
| **Cdr1as** | cicrular<br>linear | GTCTTCCAACATCTCCACAT<br>ACATCTTCCAGGAAATCAACG | TCACGATTGTCTGGAAGACC<br>GGAAGACATAGATTTTCATGAAGA |
| **Hipk3** | cicrular<br>linear | TATGTTGGTGGATCCTGTTC | ATTCACATGGATCTGAGGATAG<br>AATAACTCTGCGATCACGCA |
| **Zkscan1** | cicrular<br>linear | TCTGGATCCAGTTCAGGAGT | CAGTCATCATTCAGGCTCAG<br>CTGTCCCTATTGAGATTCCTC |

underlined – primers common to circular and linear transcripts

**Supplementary Table S2. Results of ddPCR analyses of circRNA expression levels.**

| | controls | | DM1/DM2 | | p-value (t-test) | DM1/DM2 status |
|---|---|---|---|---|---|---|
| | mean fraction | SD | mean fraction | SD | | |
| | | | **CL1_DM1_H-** | | | |
| circASXL1 | 0.10 | 0.05 | 0.11 | 0.01 | 0.91 | ↑ |
| circCAMSAP1 | 0.31 | 0.05 | 0.32 | 0.02 | 0.69 | ↑ |
| circFAM13B | 0.16 | 0.02 | 0.20 | 0.05 | 0.31 | ↑ |
| circHIPK3 | 0.41 | 0.08 | 0.45 | 0.05 | 0.48 | ↑ |
| circMBOAT2 | 0.06 | 0.02 | 0.05 | 0.01 | 0.51 | ↓ |
| circMIB1 | 0.10 | 0.03 | 0.10 | 0.01 | 1.00 | ↓ |
| circNFATC3 | 0.32 | 0.07 | 0.31 | 0.05 | 0.89 | ↓ |
| circPHC3 | 0.08 | 0.00 | 0.07 | 0.03 | 0.55 | ↓ |
| circPIP5K1C | 0.08 | 0.02 | 0.08 | 0.03 | 0.87 | ↓ |
| circSCMH1 | 0.13 | 0.02 | 0.14 | 0.01 | 0.42 | ↑ |
| circSHKBP1 | 0.03 | 0.01 | 0.03 | 0.01 | 1.00 | ↓ |
| circUBAP2_e9-12 | 0.04 | 0.01 | 0.06 | 0.02 | 0.09 | ↑ |
| circUBAP2_e7-8 | 0.15 | 0.03 | 0.17 | 0.06 | 0.66 | ↑ |
| circZKSCAN1 | 0.20 | 0.01 | 0.25 | 0.02 | 0.02 | ↑ |
| circCCDC134 | 0.05 | 0.01 | 0.06 | 0.02 | 0.55 | ↑ |
| circFOXK2 | 0.05 | 0.01 | 0.07 | 0.02 | 0.27 | ↑ |
| circPDCD11 | 0.06 | 0.00 | 0.07 | 0.02 | 0.55 | ↑ |
| circPROSC | 0.02 | 0.01 | 0.02 | 0.01 | 0.79 | ↑ |
| circCDR1as | 0.90 | 0.11 | 0.87 | 0.09 | 0.70 | ↓ |
| circMBNL1 | 0.00 | 0.00 | 0.00 | 0.00 | 0.82 | ↑ |
| | | | **CL2_DM1_H+** | | | |
| circASXL1 | 0.59 | 0.08 | 0.70 | 0.13 | 0.37 | ↑ |
| circCAMSAP1 | 0.51 | 0.06 | 0.55 | 0.05 | 0.44 | ↑ |
| circFAM13B | 0.22 | 0.13 | 0.23 | 0.12 | 0.94 | ↑ |
| circHIPK3 | 0.70 | 0.13 | 0.72 | 0.06 | 0.83 | ↑ |
| circMBOAT2 | 0.02 | 0.01 | 0.02 | 0.00 | 0.84 | ↑ |
| circMIB1 | 0.50 | 0.16 | 0.52 | 0.04 | 0.84 | ↑ |
| circNFATC3 | 0.49 | 0.07 | 0.48 | 0.08 | 0.85 | ↓ |
| circPHC3 | 0.09 | 0.08 | 0.07 | 0.02 | 0.62 | ↓ |
| circPIP5K1C | 0.08 | 0.08 | 0.03 | 0.00 | 0.26 | ↓ |
| circSCMH1 | 0.27 | 0.02 | 0.20 | 0.02 | 0.02 | ↓ |
| circSHKBP1 | 0.31 | 0.17 | 0.28 | 0.13 | 0.82 | ↓ |
| circUBAP2_e9-12 | 0.01 | 0.00 | 0.03 | 0.01 | 0.22 | ↑ |
| circUBAP2_e7-8 | 0.42 | 0.05 | 0.51 | 0.04 | 0.10 | ↑ |
| circZKSCAN1 | 0.49 | 0.12 | 0.51 | 0.14 | 0.88 | ↑ |
| circCCDC134 | 0.18 | 0.00 | 0.18 | 0.03 | 0.93 | ↑ |
| circFOXK2 | 0.33 | 0.00 | 0.27 | 0.12 | 0.54 | ↓ |
| circPDCD11 | 0.16 | 0.05 | 0.15 | 0.10 | 0.91 | ↓ |
| circPROSC | 0.03 | 0.01 | 0.02 | 0.01 | 0.33 | ↓ |
| circCDR1as | 1.00 | 0.00 | 0.96 | 0.06 | 0.50 | ↓ |
| circMBNL1 | 0.65 | 0.11 | 0.63 | 0.06 | 0.85 | ↓ |
| | | | **BP1_DM1_H-** | | | |
| circCAMSAP1 | 0.15 | 0.01 | 0.22 | 0.09 | 0.34 | ↑ |
| circFAM13B | 0.34 | 0.00 | 0.26 | 0.03 | 0.01 | ↓ |
| circHIPK3 | 0.07 | 0.02 | 0.11 | 0.06 | 0.41 | ↑ |
| circMIB1 | 0.30 | 0.03 | 0.25 | 0.06 | 0.37 | ↓ |
| circNFATC3 | 0.22 | 0.01 | 0.27 | 0.11 | 0.52 | ↑ |
| circPHC3 | 0.15 | 0.03 | 0.10 | 0.03 | 0.04 | ↓ |
| circPIP5K1C | 0.04 | 0.02 | 0.05 | 0.01 | 0.70 | ↑ |
| circSCMH1 | 0.22 | 0.06 | 0.25 | 0.07 | 0.56 | ↑ |
| circUBAP2_e9-12 | 0.04 | 0.00 | 0.05 | 0.02 | 0.61 | ↑ |
| circUBAP2_e7-8 | 0.21 | 0.00 | 0.21 | 0.07 | 0.95 | ↑ |
| circZKSCAN1 | 0.11 | 0.02 | 0.14 | 0.05 | 0.47 | ↑ |
| circFOXK2 | 0.01 | 0.00 | 0.03 | 0.03 | 0.49 | ↑ |
| circPROSC | 0.02 | 0.01 | 0.02 | 0.01 | 0.54 | ↑ |
| circCDR1as | 0.49 | 0.01 | 0.50 | 0.02 | 0.58 | ↑ |
| circMBNL1 | 0.00 | 0.00 | 0.00 | 0.00 | 0.76 | ↑ |

| | | | | | | |
|---|---|---|---|---|---|---|
| | | | **BP2_DM1_H+** | | | |
| circCAMSAP1 | 0.52 | 0.17 | 0.65 | 0.15 | 0.27 | ↑ |
| circFAM13B | 0.33 | 0.07 | 0.25 | 0.08 | 0.12 | ↓ |
| circHIPK3 | 0.26 | 0.06 | 0.36 | 0.13 | 0.12 | ↑ |
| circMIB1 | 0.74 | 0.08 | 0.73 | 0.13 | 0.91 | ↓ |
| circNFATC3 | 0.28 | 0.10 | 0.42 | 0.11 | 0.06 | ↑ |
| circPHC3 | 0.09 | 0.03 | 0.16 | 0.12 | 0.16 | ↑ |
| circPIP5K1C | 0.33 | 0.10 | 0.51 | 0.12 | 0.08 | ↑ |
| circSCMH1 | 0.53 | 0.13 | 0.61 | 0.14 | 0.42 | ↑ |
| circUBAP2 2 | 0.58 | 0.16 | 0.56 | 0.12 | 0.81 | ↓ |
| circZKSCAN1 | 0.71 | 0.07 | 0.52 | 0.13 | 0.03 | ↓ |
| circFOXK2 | 0.10 | 0.04 | 0.22 | 0.12 | 0.08 | ↑ |
| circPROSC | 0.09 | 0.03 | 0.13 | 0.08 | 0.27 | ↑ |
| circCDR1as | 0.53 | 0.04 | 0.50 | 0.03 | 0.15 | ↓ |
| | | | **BP3_DM1_H-** | | | |
| circASXL1 | 0.02 | 0.01 | 0.03 | 0.03 | 0.67 | ↑ |
| circCAMSAP1 | 0.51 | 0.15 | 0.54 | 0.34 | 0.88 | ↑ |
| circHIPK3 | 0.31 | 0.07 | 0.50 | 0.23 | 0.18 | ↑ |
| circNFATC3 | 0.39 | 0.11 | 0.49 | 0.19 | 0.48 | ↑ |
| circPHC3 | 0.15 | 0.05 | 0.27 | 0.24 | 0.33 | ↑ |
| circZKSCAN1 | 0.58 | 0.07 | 0.67 | 0.12 | 0.31 | ↑ |
| | | | **BP4_DM2_H-** | | | |
| circASXL1 | 0.02 | 0.01 | 0.01 | 0.01 | 0.15 | ↓ |
| circCAMSAP1 | 0.51 | 0.17 | 0.63 | 0.18 | 0.27 | ↑ |
| circHIPK3 | 0.31 | 0.08 | 0.42 | 0.13 | 0.14 | ↑ |
| circNFATC3 | 0.39 | 0.12 | 0.44 | 0.10 | 0.49 | ↑ |
| circPHC3 | 0.15 | 0.05 | 0.21 | 0.09 | 0.27 | ↑ |
| circZKSCAN1 | 0.58 | 0.08 | 0.63 | 0.12 | 0.40 | ↑ |
| | | | **MM_DM1_H-** | | | |
| circCAMSAP1 | 0.11 | 0.01 | 0.15 | 0.03 | 0.0009 | ↑ |
| circHIPK3 | 0.05 | 0.01 | 0.07 | 0.02 | 0.002 | ↑ |
| circZKSCAN1 | 0.007 | 0.002 | 0.009 | 0.004 | 0.2 | ↑ |
| circCDR1as | 0.65 | 0.23 | 0.50 | 0.07 | 0.07 | ↑ |

↑ - circRNA level increased; ↓ - circRNA level decreased
